# Protein•DNA mesh assembly drives dsDNA-specific and duplex length-dependent activation of cGAS

**DOI:** 10.64898/2026.05.19.726263

**Authors:** Shuai Wu, Christina M. Stallings, Stephanie M. Torres, Brendan Antiochos, Jungsan Sohn

## Abstract

Cyclic G/AMP synthase (cGAS) forms condensates on dysregulated double-stranded (ds) DNA to trigger inflammatory responses. Currently, how it specifically recognizes dsDNA and why activation depends on duplex length remain poorly understood. Using cryo-electron microscopy, biochemical assays, and single-molecule methods, we show that full-length cGAS assembles a protein•dsDNA mesh by reiteratively propagating dimers-of-dimers, driving an associative phase transition. Previously uncharacterized N- and C-terminal interactions, together with inter-dimer junction-loops, track the B-form groove and bend dsDNA to build an extensive mesh network spanning multiple duplexes. These interactions are critical for cross-stabilizing active cGAS, revealing why higher-order assembly on long duplexes maximizes signaling activity. Remarkably, cGAS can build the protein•dsDNA mesh on a single contiguous duplex by entangling its oligomerization platform, resulting in a mechanically resilient and kinetically stable signaling platform. Together, our findings establish higher-order mesh assembly of cGAS as the foundation for dsDNA selectivity, duplex-length-dependent activation, and condensate formation.

## Introduction

Phase-transitioned condensates through multivalent protein•nucleic-acid interactions are ubiquitously found in many key biological processes, ranging from the regulation of chromatin dynamics to the assembly of innate immune signaling complexes ^1–5^. Despite their prevalence, little is known about the architectural framework that underpins condensate formation at the molecular level. Moreover, why such elaborate supra-assemblies are even necessary remain often poorly understood. We report here that cyclic-G/AMP (cGAMP) Synthase (cGAS) creates a mesh-like network with double-stranded (ds)DNA, providing a structural framework that links condensate formation to nucleic acid specificity and dsDNA-length dependent activation.

Cells detect unchromatinized dsDNA as a danger arising from infection, genome instability, and dysfunctional organelles ^6–9^. Upon binding to and dimerizing on such rogue dsDNA, cGAS cyclizes ATP and GTP into cGAMP, which is the second messenger that activates stimulator of interferon genes (STING)-dependent inflammatory responses ^6,10–12^. cGAS is integral to host defense against diverse pathogens (e.g., HIV and *L. monocytogene*) ^13–15^, yet its dysregulation leads to various autoimmune and neurodegenerative diseases (e.g., Aicardi Goutières syndrome and Parkinson’s Disease) ^16–20^. Beyond innate immunity, cGAS has emerging roles in genome maintenance, such as regulating replication efficiency and chromatin dynamics, both of which are also critical for regulating tumor formation ^21–24^.

cGAS consists of an unstructured N-terminal domain (∼150 amino acids, a.a.) and a structured C-terminal catalytic domain that adopts a nucleotidyl-transferase (NTase) fold (∼350 a.a.). cGAS binds dsDNA without sequence specificity ^10–12,25–28^, and it was first thought that simply binding a DNA duplex as short as 13 base-pairs (bps) in a 1:1 stoichiometry at the catalytic domain (cGAS^cat^) is sufficient for activation ^10,28^. However, subsequent studies revealed that cGAS^cat^ forms a 2:2 dimer by sandwiching two separate dsDNA fragments ^11,12^; these dimers then further oligomerize into condensates ^29^. While earlier models described these higher-order assembles as purely liquid-liquid phase-separated condensates ^29^, more recent studies revealed that human cGAS undergoes an associative phase-transition (i.e., percolation ^30–33^), resulting in gel-like condensates in which the disordered N-terminal domain modulates material properties to prevent solidification ^34,35^.

Despite these advances, two fundamental mechanistic questions remain unresolved. First, how does cGAS achieve dsDNA-specific activation (vs. single-stranded (ss)DNA, ssRNA, or dsRNA)? Although cGAS can bind and form condensates with virtually all nucleic acids, only dsDNA can activate its enzymatic activity. We recently found that gel-like cGAS condensates are exclusively formed with cognate dsDNA ^34^. By contrast, condensates formed on other nucleic acids remain liquid-like (i.e., segregative phase separation ^30–33^), as these noncognate nucleic acids fail to support the dimerization required for activation ^34^. However, the structural basis for this selectivity is unknown.

Second, why is cGAS activation strongly dependent on duplex length? While crystallographic studies suggest that dsDNA as short as ∼15-bp can accommodate one dimer, significantly longer (≥ 45-bp) dsDNA is necessary to induce cGAS activation and oligomerization, and maximal activation occurs on ≥ 100-bp duplexes both *in vitro* and in cells (e.g., interferon (IFN) signaling by ISD45 DNA) ^25–27,29,36^. Based on the arrangement of cGAS^cat^•dsDNA complexes in the crystal lattice ^27^, currently prevailing model proposes that cGAS dimers assemble into ladder-like arrays along two parallel DNA duplexes, with increasing DNA length enhancing activity through avidity effects ^27^. Within this framework, these ladders then randomly cluster to form condensates ^29,37^. However, our previous biochemical and imaging studies suggested that cGAS assemblies might not adopt such a linear array ^26,34^. Additionally, considering that pathogenic dsDNA is often a very long single contiguous duplex (e.g., herpes virus genome, > 150-kilo-bps), how cGAS would form dimers or even higher-oligomers on such a large linear scaffold remains entirely speculative.

Here, cryo-ME structures of full-length mouse cGAS in complex with dsDNA show that a nonparallel arrangement of a dimer-of-dimers represents the minimal functional unit of cGAS, which then propagates over multiple DNA duplexes form a protein•dsDNA mesh instead of a ladder. This highly interwoven supra-structure provides a unifying framework for how cGAS interacts with dsDNA, which includes the recognition of the B-form architecture, length-dependent activation, and percolation-driven condensate formation.

## Results

### Percolation and gel-like condensate formation are preserved between human and mouse cGAS

cGAS orthologues have varying condensate forming capabilities, with the human enzyme being the most proficient due to an additional dsDNA-binding site ^37,38^. However, dimerization at the catalytic domain is essential for the dsDNA- and dsDNA-length dependent activation of all mammalian cGAS ^11,26,34,39^. Moreover, the gel-like characteristics of hcGAS•dsDNA condensates arise from dimerization, not the additional binding site ^34^. Thus, we first asked if dimerization drives the percolation of cGAS in a distant species. We employed full-length mouse (m)cGAS^(FL)^, as it has the weakest condensate forming capability in mammals ^38^. The mouse enzyme formed droplet-like condensates with fluor-labeled 60-bp dsDNA (dsDNA_60_), while those formed with 100-bp dsDNA showed a mixture of round and mesh-like morphologies; 300-bp duplexes produced even more extended mesh-like condensates (Figures 1A and S1A-B). These observations are consistent with the dsDNA length (dimerization)-dependent condensation of hcGAS^FL^ ^34,35^. Tracking fluorescence recovery after photobleaching (FRAP) indicated that even condensates formed with 60-bp duplexes have gel-like characteristics (t_1/2_ > 10 min ^34^), which in turn suggest that cGAS condensates are formed by coupling associative and segregative phase transitions ^30–33^ (Figures 1B-C and S1C). The intra-condensate dynamics were inversely correlated with dsDNA lengths (Figures 1B-C and S1C), indicating that percolation dominates with the increased multivalency of longer duplexes ^5,30–33^.

**Figure 1:**
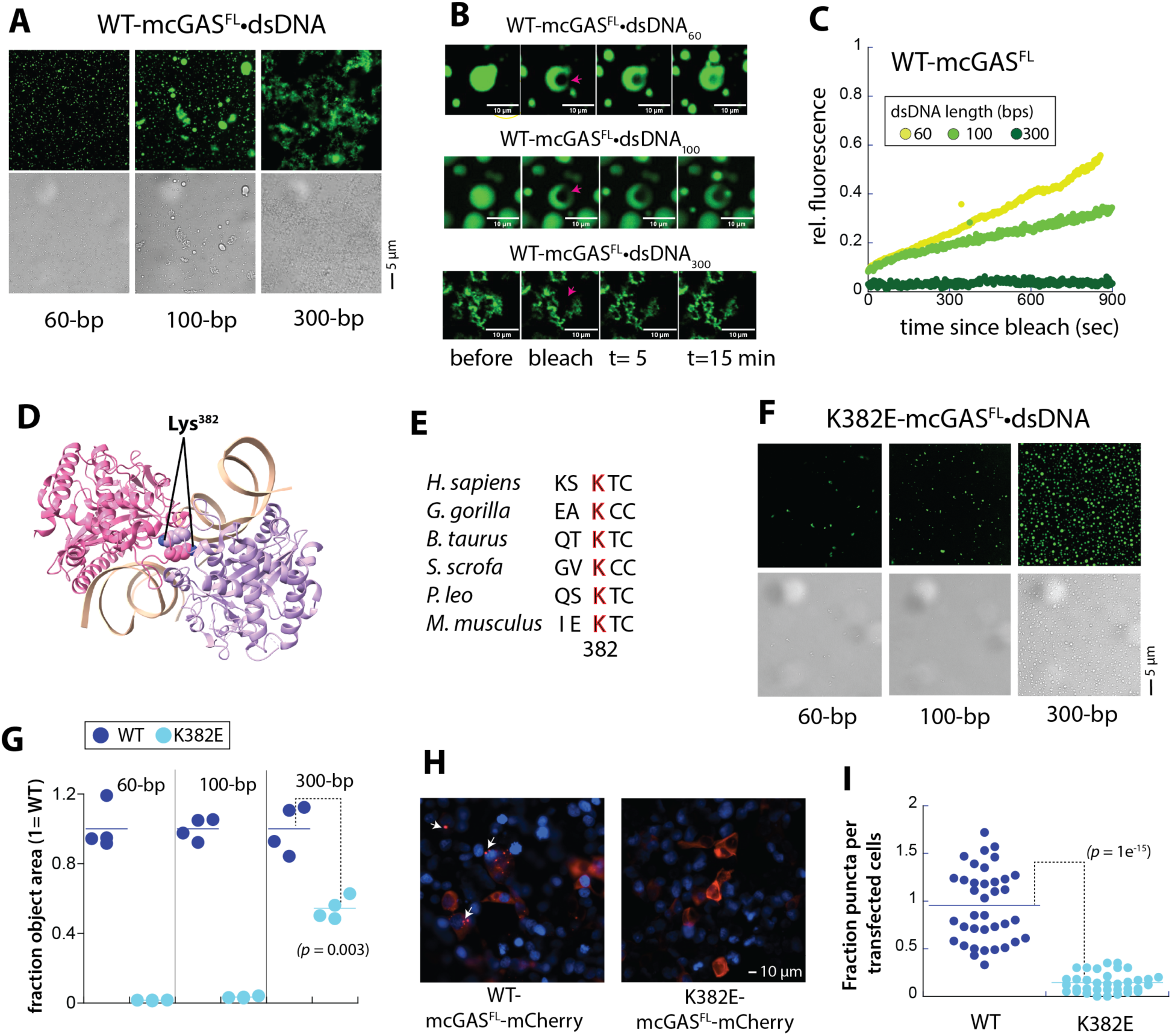
Gel-like condensate formation is conserved in mouse cGAS. **(A)** Fluorescent and bright-field images of WT-mcGAS^FL^ condensates formed with fluor-labeled dsDNA of indicated lengths (3 µM for both protein and dsDNA, *t* = 25 min; fluorescein amidte (FAM) for 60-bp and 100-bp, and Alexa_488_ for 300-bp dsDNA). **(B)** Representative FRAP images of WT-mcGAS^FL^ condensates formed with the fluor-labeled dsDNA of indicated lengths. **(C)** A representative plot showing the recovery of fluorescence over time after bleaching from Figure 1B. **(D)** A crystal structure of the mcGAS^cat^ dimer bound to dsDNA (PDB: 7UUX). Each monomer is colored differently and the conserved lysine at the dimer interface is shown as spheres. **(E)** The a.a. sequence alignment at the dimerization loop region showing the conserved Lys. **(F)** Fluorescent and bright-field images of cGAS condensates formed on fluor-labeled dsDNA of indicated lengths (3 µM for both protein and dsDNA, *t* = 30 min). **(G)** Plots showing relative amounts of condensates (area covered) formed by each cGAS construct with indicated dsDNA after incubating for 5 minutes. Data were normalized to the number of condensates observed for WT. Each dot represents one image (view field). All *p* values are calculated using a two-tailed *t*-test assuming unequal variance. **(H)** Images showing mCherry-tagged WT- or K382E-mcGAS^FL^ condensates in HEK293T cells. White arrows indicate cGAS puncta/condensates. Blue: DAPI. **(I)** The relative number of puncta observed from HEK293T cells transfected with mCherry-tagged WT- vs. K382E-mcGAS^FL^.

Noncognate nucleic-acids such as ssRNA can dissolve human (h)cGAS•dsDNA hydrogels into liquid-like droplets by inhibiting dimerization ^34^ (i.e., decoupling the associative nature from the segregative aspect). Here, adding 60-base ssRNA disrupted the mesh-like network of mcGAS^FL^•dsDNA_300_ (Figure S1D). Moreover, ssRNA_60_ converted mcGAS^FL^•dsDNA_100_ complexes into more droplet-like entities, eventually displacing bound dsDNA_100_ by mass action (Figure S1E). Additional NaCl also disentangled cGAS•dsDNA mesh-like condensates into individual droplets (Figure S1F). Next, to test whether dimerization is also critical for the percolation of the mouse enzyme, we mutated a conserved Lys at the dimer interface (K382E; Figure 1D-E; see also ref. ^11,12,26^). The resulting mutant rarely produced condensates with 60- or 100-bp dsDNA (Figures 1F-G and S1G). With 300-bp dsDNA, K382E-mcGAS^FL^ formed round droplets even after an hour without showing the mesh-like morphology of WT (Figure 1A vs. 1F, and see also Figure S1G). Moreover, when transfected into HEK293T cells, mCherry-tagged WT formed numerous puncta (Figure 1H-I), indicative of condensate formation ^29,34,35,38^. By contrast, mCherry-tagged K382E-mcGAS^FL^ remained largely diffused (Figure 1H-I), suggesting that dimerization is even more critical for the mouse enzyme to form condensates in cells than the human ortholog ^34^. We thus surmise that dimerization leads to percolation mammalian cGAS.

### Dimers-of-dimers propagate across DNA duplexes to form a mesh-like network

Unlike segregative phase-separation, condensate formation via percolation entails iteratively establishing a network of molecular interactions ^5,30–33^. To elucidate such a framework, we examined mcGAS^FL^ bound to 66- or 100-bp dsDNA using cryo-EM. We chose these two lengths because they would be more conducive to high-resolution cryo-EM studies than the branched structures produced by 300-bp duplexes. Considering the changes in shapes and dynamics (Figures. 1A-C), we also reasoned that 100-bp dsDNA might reveal additional multivalent interactions missing from the shorter duplex (Figures. 1A-C). The N-terminal maltose binding protein (MBP) purification tag was retained in our cryo-EM samples, as we reasoned that it might facilitate particle picking (Figure S2A; MBP does not interfere with the dsDNA length-dependent activation or gel-like condensate formation of mcGAS^FL^ (Figure S1H-I)). 2D-classification showed that most particles represent various views of two cGAS dimers arranged in a head-to-head manner on dsDNA (Figures 2A, S2B and S3B). MBP was not visible (Figures 2A-B, S2B, S3B, and S4B), and the N-domain remained largely disordered, indicating that the catalytic domain dominates stable dsDNA•protein interactions. Although both 66-bp and 100-bp fragments are long enough to accommodate the ladder-like arrangement beyond the dimer of dimers ^27^, we did not find any particles consistent with such a linear configuration (a footprint of one catalytic domain is ∼15-bp ^27^; Figures S2B, S3B, and S4B). These observations suggest that a dimer of dimers constitutes the minimal functional unit of cGAS in solution (i.e., why at least ∼45-bp dsDNA such as ISD45 is necessary to activate cGAS in cells) ^25,27,36,39^.

**Figure 2:**
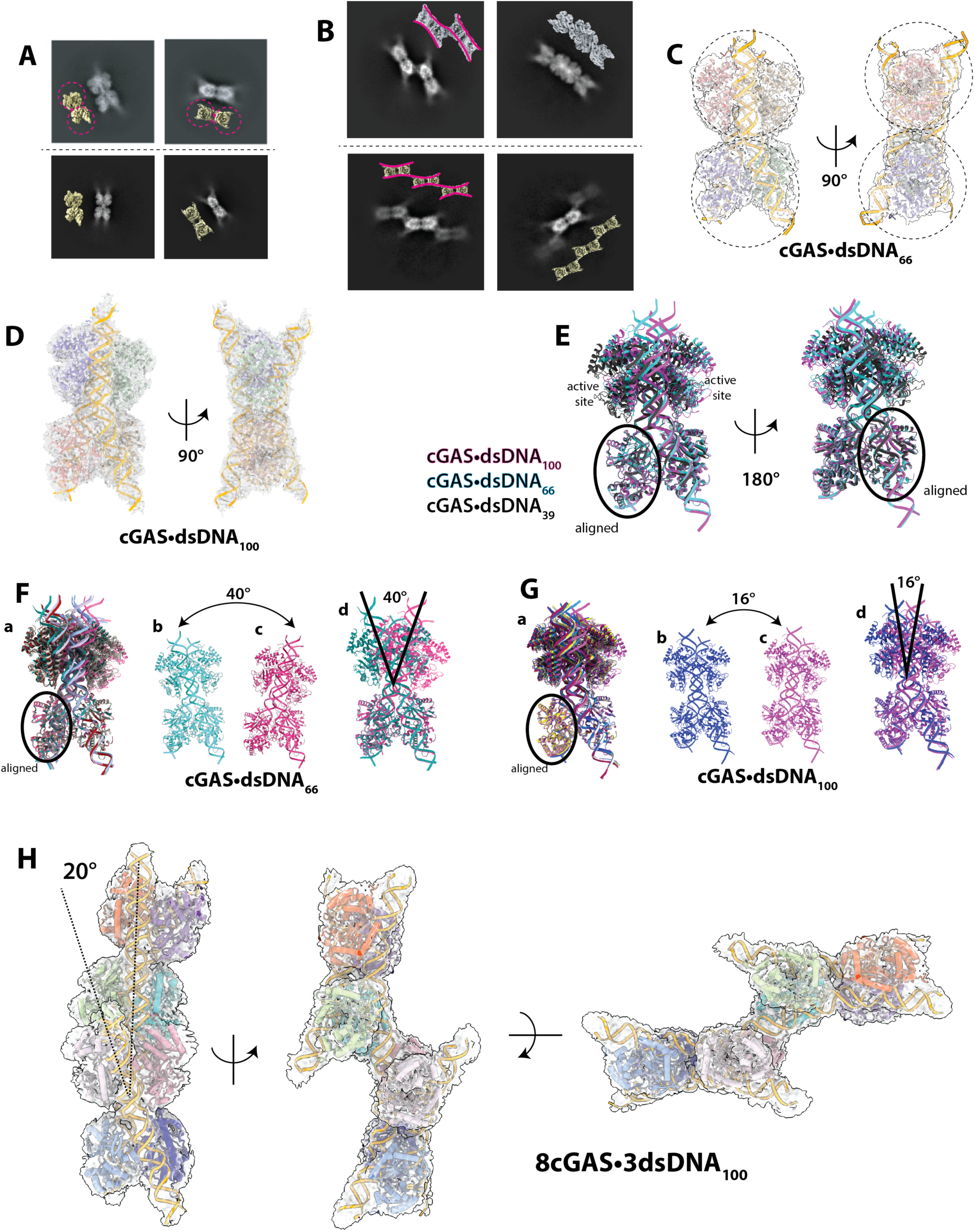
Cryo-EM structures of mcGAS^FL^ bound to 66- or 100-bp dsDNA. **(A)** 2D class averages and corresponding 3D volumes of a dimer of dimers formed by mcGAS^FL^•dsDNA_66_ (top) and mcGAS^FL^•dsDNA_100_ (bottom). Each dotted circle indicates one dimer. **(B)** 2D class averages and corresponding 3D volumes of the higher-order oligomers formed by mcGAS^FL^•dsDNA_100_. Top: two different views of a dimer of pseudo tetramers assembled on three DNA duplexes. Bottom: A trimer of pseudo-tetramers assembled on four duplexes. Each magenta line on volume maps indicates a DNA duplex (3 and 4 lines for top and bottom, respectively). **(C)** The composite cryo-EM map fit with the corresponding model of the mcGAS^FL^•dsDNA_66_ pseudo-tetramer complex. Each monomer is colored differently, and dsDNA is modeled up 50 bps. Each dotted circle indicates one dimer. **(D)** Composite cryo-EM map and model of the mcGAS^FL^•dsDNA_100_ pseudo-tetramer complex. Each monomer is colored differently, and dsDNA is modeled up 50 bps. **(E)** Structural overlay of the cryo-EM models with the crystal structure of 39-bp dsDNA bound to mcGAS^cat^ (PDB: 5N6I). **(F)** Six mcGAS^FL^•dsDNA_66_ models from the 3DVA cluster mode are aligned at one monomer (**a**); two outputs that show the biggest deviation from each other (**b**-**c**); the two most deviated models are then aligned at one monomer (**d**). **(G)** Six mcGAS^FL^•dsDNA_100_ models from the 3DVA cluster mode are aligned at one monomer (**a**); two outputs that show the biggest deviation from each other (**b**-**c**); the two most deviated models are then aligned at one monomer (**d**). **(H)** Composite cryo-EM map and model of the mcGAS^FL^•dsDNA_100_ complex composed of two pseudo-tetramers assembled on three duplexes. The modeled shared dsDNA is 84 bps and the peripheral duplexes are modeled to 50-bp. Monomers are individually colored.

We also found several 2D classes where these dimers-of-dimers (interchangeably referred as “pseudo-tetramers” hereafter) propagate over multiple 100-bp duplexes (Figure 2B). Notably, a number of particles appeared to represent various views of two cGAS dimers-of-dimers arranged diagonally over three DNA duplexes (Figures 2B top and S4B). Moreover, a subset of class averages showed three pseudo-tetramers assembled over four duplexes (Figure 2B bottom). Thus, our observations reveal that, instead of forming a linear protein•dsDNA ladder ^27^, cGAS constructs a protein•dsDNA mesh by iteratively assembling dimers-of-dimers across multiple duplexes.

### cGAS bends and twists dsDNA within and between pseudo-tetramers to build a mesh

After multiple rounds of heterogeneous and focused refinements (Figures S2 and S3, and Table S1), we obtained composite cryo-EM maps corresponding to a cGAS pseudo-tetramer bound to 66-bp or 100-bp dsDNA (2.7 Å and 2.9 Å resolution, respectively; Figures 2C-D and S5A). In comparison to the crystal structure of two mcGAS^cat^ dimers bound to two 39-bp duplexes ^27^, protein•dsDNA interactions at each protomer are largely conserved in our cryo-EM structures (Figure 2E), and the two dsDNA fragments were also concaved in the same manner between to dimers (Figure 2C-E). However, aligning these structures at one monomer showed that the position of the second dimer largely deviates from one another (Figure 2E), suggesting that the two dimers sample different conformations instead of being affixed into the ladder-like parallel arrangement seen in the crystal lattice ^27^. To further probe this conformational heterogeneity, we conducted the 3D variability analysis (3DVA) in cryoSPARC. The latent coordinates revealed in the simple mode showed that the two cGAS dimers twist and turn respect to one another using the unbound dsDNA at the junction as a flexible hinge (Supplementary Movies 1-2). We next generated 6 models from each dsDNA-bound complex using the 3DVA cluster mode (Figures S2B and S3B). Aligning these structures at one monomer showed that the bound dsDNA can be bent as much as 40° and 16° for 66-bp and 100-bp dsDNA bound complexes, respectively (Figure 2F-G).

We also obtained a composite map of two cGAS pseudo-tetramers bound across three 100-bp DNA duplexes at 3.7 Å resolution (Figures 2H S4, and Table S1). The resulting model shows that each pseudo-tetramer binds dsDNA identically as the two separate pseudo-tetramers (Figures 2H and S5A-B, and Supplementary Movie 3). However, none of the three duplexes are arranged in parallel. Instead, the shared dsDNA is curved like the *tilde* symbol, thereby positioning the two pseudo-tetramers in a staggered configuration (Figures 2H and S5B-C, and Supplementary Movie 3). Our observations suggest that cGAS bends and twists the dsDNA scaffold both within and between pseudo-tetramers to form the protein•DNA mesh.

### Mesh construction tracks the B-form helix and is directly tied to the DNA duplex-length dependent activation

We noted that the loops at the junction between the two dimers within one pseudo-tetramer make variable contacts with dsDNA as they twist and turn in our 3DVA, seemingly counter-stabilizing the dsDNA-bound complex (Figures 3A and S5D, and Supplementary Movies 1-2 and 4). These junction-loops remained poorly resolved in the crystal structures of cGAS dimers, as bound dsDNA is too short to reach this region (13 to 18-bp ^10–12,28,40^) vs. 50-bp duplexes modeled in our structures. Notably, it appears that the basic residues in this loop (Lys^2^^40^, Arg^222^, and Arg^244^) hold the backbone frame while Arg^241^ senses the major groove without making any specific contacts (Figure 3A). These interactions are mirrored diagonally by another protomer from the opposing dimer, thus apparently counter-stabilizing the pseudo-tetramer (Figures 3A and S5D-E; Supplementary Movies 1-2 and 4).

**Figure 3:**
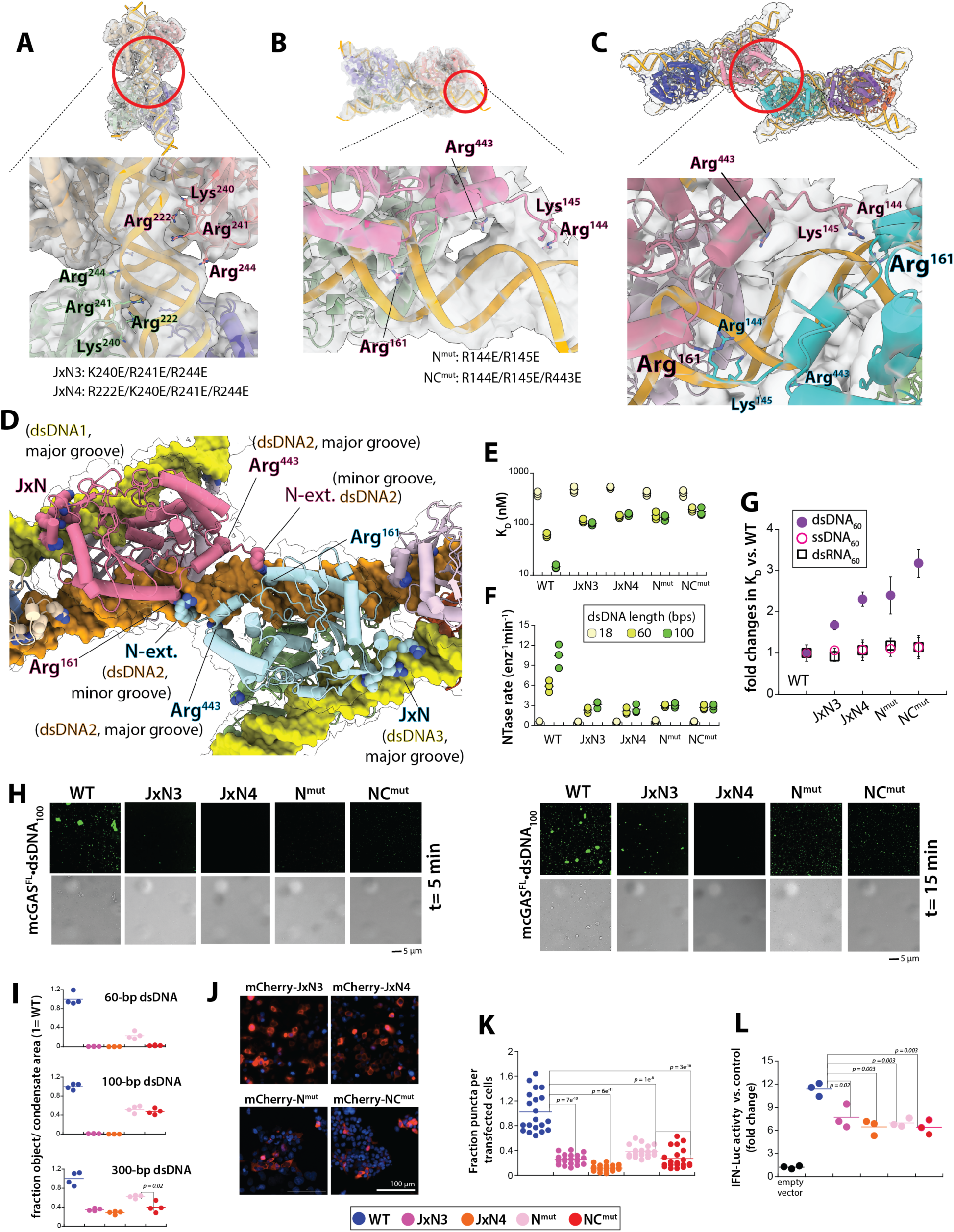
DNA groove-tracking interactions underpin condensate formation and enzymatic activation. **(A)** Composite map of a mcGAS^FL^•dsDNA_100_ pseudo-tetramer (top) and zoomed-in view (bottom) showing the junction between two dimers. Mutated residues and their IDs are indicated. **(B)** Composite map and model of a mcGAS^FL^•dsDNA_100_ pseudo-tetramer (top) and a zoomed-in view (bottom) showing contacts between dsDNA and the N-terminal extension (Arg^144^ and Lys^145^) and the C-terminal Arg^443^; Arg^161^ is also shown. Mutated residues and their IDs are indicated. **(C)** Composite map and model of a mcGAS^FL^•dsDNA_100_ dimer of pseudo-tetramers bound to three separate DNA duplexes (top) and a zoomed-in view (bottom) showing Arg^144^, Lys^145^, and Arg^443^ from two different monomers contacting dsDNA near Arg^161^ via the shared dsDNA. **(D)** Model of the mcGAS^FL^ multimer assembled over three 100-bp DNA duplexes highlighting the DNA groove tracking interactions. The interactions that track either the major or minor groove of each dsDNA (dsDNA1-3) from two different monomers (pink and cyan) are shown as the sphere representation: N-terminal extension (N-ext.), Arg^443^ from the C-terminus, and the junction region (JxN). **(E)** A plot showing the binding affinity (K_D_) of each mcGAS^FL^ construct against dsDNA with indicated length. **(F)** A plot showing the catalytic (NTase) activity of each mcGAS^FL^ construct against ATP:GTP (1:1) and indicated dsDNA. **(G)** A plot showing the relative binding affinity of each mutant against indicated nucleic acids when normalized to the K_D_ of WT. **(H)** Fluorescent and bright-field images of mcGAS^FL^ condensates formed on FAM-labeled 100-bp dsDNA after incubating 5 and 15 min (3 µM for both protein and dsDNA). **(I**) Plots showing relative amounts of condensates (area covered) formed by each cGAS construct on indicated dsDNA length after incubating for 5 min. **(J)** Images showing mCherry-tagged mcGAS^FL^ mutants expressed in HEK293T cells. Blue: DAPI. **(K)** The relative number of puncta observed from HEK293T cells transfected with indicated mCherry-tagged mcGAS^FL^ variants. **(L)** Fold induction of IFN-luciferase in HEK293T co-transfected with plasmids encoding STING, IFN-Luc, and each mcGAS variant.

The disordered N-domain of cGAS (149 a.a. for mouse) has multiple regulatory roles: assisting dsDNA binding ^34,41^, enhancing dimerization ^26^, promoting condensate formation ^29,35^, and keeping the gel-like condensate from transforming into solid-like aggregates ^35^. Although remained mostly invisible, we noted the N-terminal extension (especially Arg^144^ and Lys^145^) interacting with the dsDNA backbone frame of a minor groove in both 66- and 100-bp bound pseudo-tetramer structures (Figures 3B and S5D-E). Moreover, Arg^443^ from the C-terminus contacts the backbone phosphates of the adjacent major groove (Figures 3B and S5D-E). All previously reported cGAS structures failed to reveal these interactions, because either the employed catalytic domain (a.a., 147-507 in mouse) or dsDNA fragments were too short (≤ 39-bp) ^10,11,27,28^.

In our 3DVA, it appears that the N-terminal extensions coordinate with the junction loops to cross-stabilize the pseudo-tetramer (Supplementary Movies 1-2 and 4). More importantly, these extended interactions occur immediately upstream of the “spine helix” that “breaks” upon dsDNA-binding to initiate the allosteric activation within each protomer (Figures 3B and S5E-F) ^5,10,12,28^. That is, Arg^161^ (Arg^176^ in human) in the spine helix flips up upon binding dsDNA to make the only side-chain•base interaction in the complex to allosterically restructure the active site (Figure S5F; mutating this residue abrogates dsDNA-dependent activation ^10,11,28,39^). Thus, our structures suggest that the N-terminal extension augments the dsDNA-dependent activation of cGAS *in cis*. Furthermore, in the higher-order structure, these interactions extend across multiple pseudo-tetramers: the N-terminal residues of a protomer from one pseudo-tetramer bind the same minor groove that Arg^161^ of another protomer from a different pseudo-tetramer penetrates (Figure 3C, pink N-domain residues to cyan Arg^161^, and *vice versa*; see also Figure 3D). Likewise, Arg^443^ from one pseudo-tetramer binds the major groove directly upstream to the minor groove that Arg^161^ from the diagonal pseudo-tetramer penetrates (Figure 3C-D). Thus, our observations suggest that mesh formation enables cGAS to self-reinforce dsDNA-binding and dsDNA-dependent activation both *in cis* and *in trans*.

Our structures shed light into several long-standing questions regarding the activation mechanism of cGAS. First, protein•dsDNA interactions at the N/C-terminal extension and the junction put the footprint of one full-length cGAS to at least ∼22-bp (e.g., Figure S5E), explaining why short duplexes used in crystallographic studies are especially ineffective in binding and activating cGAS (13 to 18-bp) ^25,27,39,40^. Second, the non-parallel arrangements of cGAS dimers within and between pseudo-tetramers suggest that the junction interactions that apparently bend and twist the concaved dsDNA are important not only for forming one cGAS pseudo-tetramer (minimal functional unit), but also for constructing the mesh (Figures 2F and 3A). Third, the *in-trans* interactions across the shared dsDNA by the N- and C-terminal residues show why higher-order assembly on “long” dsDNA is necessary for optimal activation (Figures 2H and 3C-D). Finally, all these interactions track either the major or minor groove of multiple duplexes (Figures 3D and S5D-E), suggesting that the dsDNA-dependent activation, thus mesh construction, is accomplished by recognizing the B-form architecture.

### Biochemical and cellular experiments support our cryo-EM structures

To test our structural observations, we made two combinatory charge-reversal mutations on the junction loops (R222E/K240E/R241E: termed JxN3, R222E/K240E/R241E/R244E: termed JxN4; Figure 3A and D). We also mutated the two positively charged residues at the N-terminal extension with or without R443E (R144E/K145E: termed N^mut^, R144E/K145E/R443E: termed NC^mut^; Figure 3B-D). All four recombinant mcGAS^FL^ mutants bound FAM-labeled 18-bp dsDNA as tightly as WT (< 1.3-fold difference), suggesting that the original residues contribute minimally in forming one dimer and dsDNA_18_ is too short to reach these regions (Figure 3E). However, the strong dependence on dsDNA-length for both binding and activation seen from WT was greatly diminished in all four mutants. For instance, compared to 18-bp dsDNA, the binding affinity of WT improved 6.4- and 27-fold for 60- and 100-bp dsDNA, respectively (Figure 3E). By contrast, the mutants bound 60-bp dsDNA only 2.1 to 4.6-fold more tightly than the 18-bp duplex (Figure 3E). Moreover, all four mutants bound 100-bp dsDNA with essentially the same affinity as the 60-bp duplex, resulting in the ∼10-fold weaker affinity vs. WT (dark green in Figure 3E). Although the N-domain seemed to be more critical in binding the longer duplexes than Arg^443^ (Figure 3E), the R443E mutant bound both 60- and 100-bp dsDNA ∼2-fold more weakly than WT (Figure S5G), supporting its role in mesh construction. The catalytic activities of mutants mirrored these trends: compared to 18-bp dsDNA, the enzymatic NTase activity ^26,39,40^ of WT against a 1:1 mixture of ATP and GTP improved 9- and 17-fold for 60- and 100-bp duplexes, respectively (Figure 3F). By contrast, the NTase activity of all four mutants increased only ∼4-fold between 18-bp and 60-bp dsDNA; again with no further enhancement with 100-bp dsDNA (Figure 3F). These results demonstrate that the observed interactions in our cryo-EM structures are critical for dsDNA length-dependent binding and activation of cGAS. Additionally, unlike binding 60-bp dsDNA, the binding affinities of WT and all four mutants did not change toward 60-bp dsRNA or 60-base ssDNA, supporting the mechanism in which these residues are exclusively involved in recognizing dsDNA (Figure 3G).

To test their roles in condensate formation, we imaged the mutant proteins in complex with 60-, 100-, and 300-bp dsDNA (Figures 3H and S6). The ability to form condensates on 60-bp dsDNA was abrogated for JxN3 and JxN4 (Figures 3I and S6A-B). With 100-bp dsDNA, JxN3 produced only a few condensates while JxN4 still failed (Figures 3H-I and S6A-B). Although both mutants showed numerous condensates with 300-bp dsDNA (Figure S6A-B), they formed slower (Figure 3I) and remain droplet-like without showing the mesh-like morphology of WT (Figure S1B vs. S6A-B). These results suggest that junction-loop interactions are essential not only for stabilizing one pseudo-tetramer, but also for propagating them for mesh assembly.

N^mut^ and NC^mut^ exhibited similar defects. With 60-bp dsDNA, both mutants formed fewer condensates and did so more slowly than WT (Figures 3I and S6C-D). With 100-bp dsDNA, they produced smaller isolated condensates without forming mesh-like extended oligomers (Figures 3H-I and 6C-D). Although 300-bp dsDNA supported the formation of mesh-like entities for N^mut^ and NC^mut^, they formed more slowly (Figure 3I) and appeared less extended than WT (Figure S6C-D). Additionally, NC^mut^ produced condensates more slowly than N^mut^ on 300-bp dsDNA (Figure 3I), again indicating that Arg^443^ supports mesh construction (Figure S6C-D). When transfected into HEK293T cells as mCherry-tagged variants, all mutants remain largely diffused and showed significantly fewer puncta than WT (Figure 1H vs. 3J-K). Further supporting that mesh construction is integral to the signaling activity of cGAS, upon co-transfecting into HEK293T cells with plasmids encoding STING and the IFN-luc reporter, all mutants showed significantly decreased luciferase activities than WT (Figure 3L). All our results consistently support both our cryo-EM structures and the signaling mechanism in which mesh construction is integral to both the dsDNA-specific and dsDNA length-dependent activation of cGAS.

### Mesh construction on dsDNA by cGAS results in kinetically stable and mechanically resilient signaling complexes

cGAMP synthesis occurs in minutes ^10,26,40^, which is rather slow for a signaling enzyme. Thus, we speculated that one of the reasons why cGAS forms a percolated mesh in a dsDNA length-dependent manner is to prolong the lifetime of the active signaling complex, as the length of duplex is directly correlated with the severity of intracellular maladies ^25,27^ (e.g., intact viral genome vs. degraded fragments). To test this idea, we compared the off-rate kinetics of bound dsDNA in various lengths from WT vs. the mutants impaired in mesh construction (as shown in Figure 4A: K382E (dimerization), JxN3 and JxN4 (stabilizing one pseudo-tetramer), and N^mut^ and NC^mut^ (stabilizing pseudo-tetramers and propagating mesh). For WT, the half-life of bound complex increased ∼10-fold between 24- and 60-bp (Figure 4B; ∼15 vs. ∼120 sec; see also Figure S7A), and less than 40% of FAM-labeled 100-bp dsDNA dissociated (Figure 4C and the plateaus in Figure S7A); the lack of complete competition is likely due to the gel-like nature/co-condensing behaviors ^34^. By contrast, all mutants failed to show appreciable increase in dsDNA half-life with the longer duplexes, and at least 60% of the bound 100-bp dsDNA dissociated (Figures 4B-C and S7A). These observations suggest that mesh construction is critical for the kinetic stability of the active cGAS•dsDNA complex.

**Figure 4:**
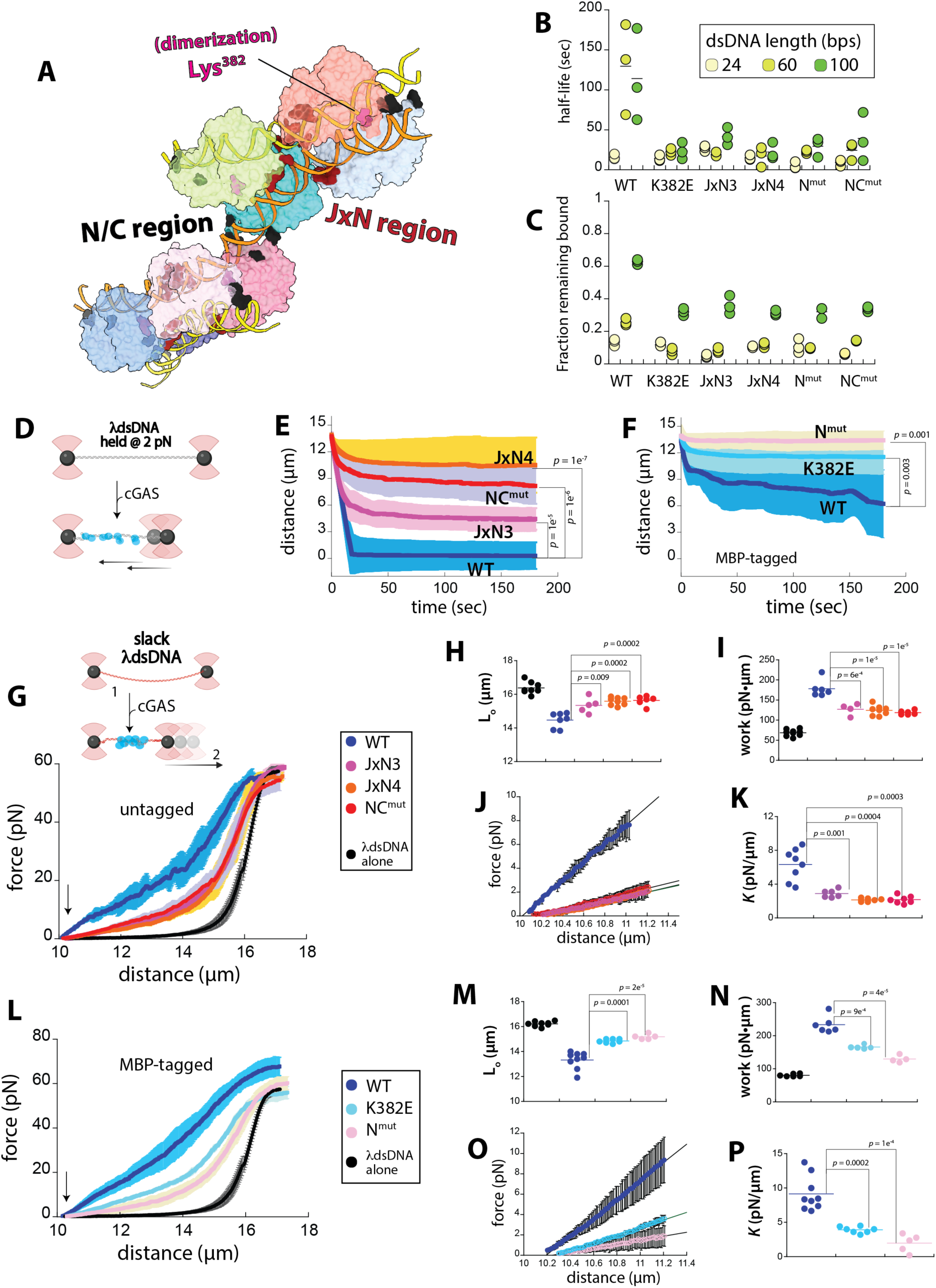
Mesh construction underpins the assembly of a robust cGAS•dsDNA signaling complex that is both kinetically stable and mechanically resilient. **(A)** A cartoon showing two cGAS pseudo-tetramers assembled on three 100-bp DNA duplexes. Lys^382^, the JxN region (Arg^222^, Lys^240^, Arg^241^, Arg^244^) and the N/C-term region (Lys^144^, Arg^145^, and Arg^443^) are colored and labeled. **(B)** A plot showing the lifetime (half-life) of indicated cGAS construct in complex with FAM-labeled 24-, 60-, or 100-bp dsDNA upon introducing excess unlabeled dsDNA_60_ (see also Figure S7A) **(C)** A plot showing the fraction of each mcGAS^FL^ construct remaining bound to different lengths of FAM-labeled dsDNA after adding excess unlabeled dsDNA_60_ (i.e., the plateau of each plot shown in Figure S7A). **(D)** A schematic of the force clamp (FC) experiment using the C-Trap. **(E)** A plot showing the average distance between two traps over time upon adding each untagged mcGAS^FL^ construct (3 µM). *n* ≥ 5 ± standard deviations (shaded area). **(F)** A plot showing the average distance between two traps over time upon adding each MBP-tagged mcGAS^FL^ (3 µM). *n* ≥ 5 ± standard deviations (shaded area). **(G)** Top: A schematic of the force extension (FE) experiment using the C-Trap. Bottom: A plot showing the average force generated vs. distance upon pulling κdsDNA in complex with each untagged mcGAS^FL^ (3 µM). *n* ≥ 5 ± standard deviations (shaded areas). **(H)** A plot showing the estimated contour length (L_0_) of κdsDNA alone or when in complex with indicated mcGAS^FL^ (i.e. pulled distance at 30 pN; see also Figure S7B for an extensible worm-like-chain model for naked κdsDNA, which is consistent with the L_(x)_ at 30 pN ≈ L_0_) ^42^. **(I)** A plot showing the work performed to stretch either κdsDNA alone or when bound to untagged-mcGAS^FL^ proteins. **(J)** A plot of average force generated whiling stretching the first ∼ 1 µm. Error bars are determined by standard deviations as seen in Figure 4G. Lines are linear fits indicative of stiffness (force/distance). **(K)** A plot showing the stiffness (*K*) of WT or mutant bound κdsDNA as estimated by the slope of each FE curve while pulling the first ∼ 1 µm. **(L)** A plot showing the average force generated vs. distance upon pulling κdsDNA in complex with each MBP-tagged mcGAS^FL^ (3 µM). *n* ≥ 5 ± standard deviations (shaded areas). **(M)** A plot showing the estimated contour length (L_0_) of κdsDNA alone or when in complex with indicated MBP-mcGAS^FL^ (i.e. pulled distance at 30 pN as described in 4H) **(N)** A plot showing the work performed while pulling either κdsDNA alone or when bound to each MBP-mcGAS^FL^ protein. **(O)** A plot of average force generated whiling stretching the first ∼ 1 µm. Error bars are determined by standard deviations as seen in Figs. 4G and L. Lines are linear fits indicative of stiffness (force/distance). **(P)** A plot showing the stiffness (*K*) of WT or mutant bound κdsDNA as estimated by the slope of each FE curve while pulling the first ∼ 1 µm.

It is noteworthy that pathogenic dsDNA such as viral genome often presents itself as one single duplex ranges in kilo-bps (e.g., hepres virus genome is ∼150-kilo bps). Thus, for cGAS to form a mesh-like network with such one contiguous dsDNA, it must be able to mechanically compact its oligomerization platform. To test this idea, using an optical trap, we tracked whether cGAS can shrink κdsDNA held at 2 pN (Figure 4D; i.e., force-clamp (FC)). WT completely contracted κdsDNA, demonstrating that cGAS can mechanically manipulate dsDNA (Figure 4E; ∼16 µm). By contrast, JxN3, JxN4, and NC^mut^ compacted κdsDNA only between 4 to 10 µm, despite present at saturating concentrations (Figure 4E; 3 µM, ∼30-fold higher than K_D_ for 100-bp dsDNA). K382E (K382A) and N^mut^ mutants repeatedly broke tethered κdsDNA during our single-molecule experiments for inexplicable reasons (not shown), but retaining the N-terminal MBP tag circumvented this issue. _(MBP)_WT compacted κdsDNA nearly 8 µm, while _(MBP)_K382E contracted only ∼ 2 µm; _(MBP)_ N^mut^ failed to compact κdsDNA altogether (Figure 4F). Our results suggest that cGAS can bend and entangle one long DNA duplex to build the mesh.

We then conducted force-extension (FE) experiments in which we recorded force generated while stretching κdsDNA bound to each cGAS protein (Figure 4G top). cGAS-bound complexes immediately required more force to stretch than naked κdsDNA (Figure 4G; colored vs. black, indicated by an arrow). Moreover, comparing the pulled distance at 30 pN ^42^ suggest that the contour (L_0_, overall) lengths of cGAS•κdsDNA complexes are at least 1 µm shorter than unbound κdsDNA (Figure 4H; one cGAS is ∼ 6 nm long; see also Figure S7B). Our observations consistently suggest that cGAS generates condensation force by entangling dsDNA ^43–47^. Moreover, compared to κdsDNA bound to each mutant, not only is the contour length of the WT-bound complex significantly shorter (Figure 4H), but it also requires more work to stretch (Figure 4I). Comparing the slopes of FE curves for the first ∼ 1 µm indicated that the WT•κdsDNA complex is significantly stiffer than those bound to mutants as we initiate each pull (Figure 4J-K).

Likewise, pulling κdsDNA bound to each MBP-tagged cGAS variant generated more force than stretching naked κdsDNA (Figure 4L, indicated by an arrow) and resulted in shorter contour lengths (Figure 4M). Significantly more work was also performed while pulling the WT-bound complex than stretching κdsDNA bound to either K382E or N^mut^ (Figure 4N). Comparing force generated while pulling the first ∼1 µm also indicated that the _(MBP)_WT-bound complex is stiffer (Figure 4O-P). Overall, our results suggest that the mesh construction of cGAS is a force-dependent process in which cGAS builds a protein•dsDNA mesh to create a robust signaling platform that is both kinetically stable and mechanically resilient.

## Discussion

cGAS is central to the host innate defense against unchromatinized dsDNA, a major danger signal arising from various intracellular catastrophes ranging from pathogen invasion to dysfunctional mitochondria ^6–9^. We show here that the assembly of a mesh-like supra-structure is at the crux of the interaction between cGAS and dsDNA, governing nucleic-acid specificity, duplex-length dependent activation, condensation, and even mechanical transaction with dsDNA (Figure 5). Given that both human and mouse cGAS undergo DNA duplex-length dependent percolation via dimerization (Figure 1), although subtle mechanistic differences would exist, we propose that cross-stabilizing active protomers through a mesh-like network on dsDNA is central to cGAS activation across different species.

**Figure 5:**
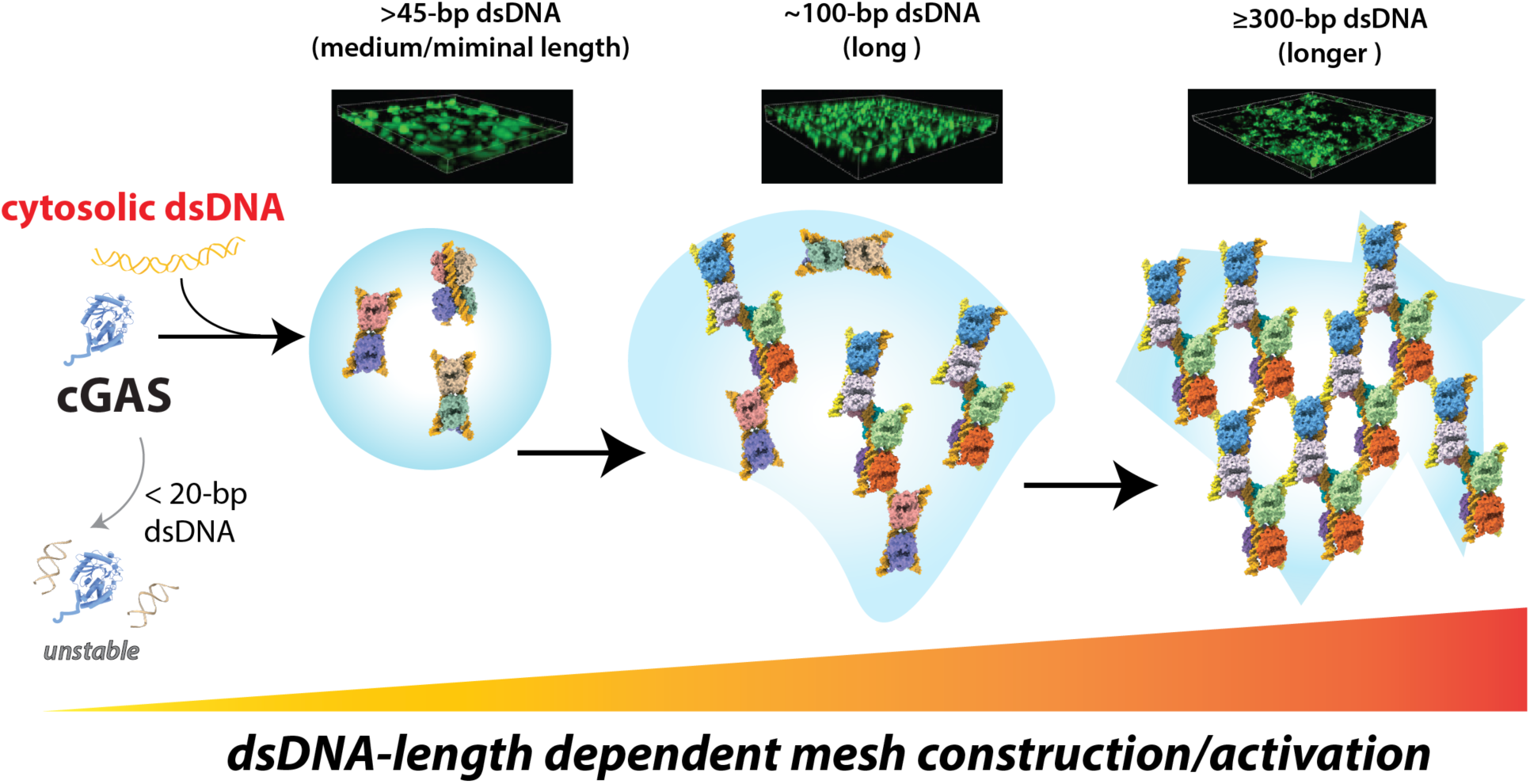
A model for dsDNA length-dependent mesh construction and activation of cGAS. Short dsDNA fragments (e.g., < 20-bp) cannot support the formation of stable cGAS dimers. dsDNA ≥ 45-bp leads to the formation of a stable dimer of dimers, which would also facilitate condensate formation by coupling segregative and associative phase transitions. With longer dsDNA (e.g,100-bp), these dimers of dimers would branch out to multiple duplexes, initiating mesh construction (percolation now becomes the more dominant organizing principle). These multimeric interactions would also allow nearby cGAS molecules to counter-stabilize their active conformations. With even longer dsDNA (e.g., 300-bp), the protein•dsDNA mesh would expand to further bolster the active conformation. cGAS would also bend and entangle such longer duplexes to facilitate mesh construction.

### The dsDNA specificity of cGAS

Early studies proposed cGAS recognizes dsDNA via a Zn^2+^-finger in each monomer ^28^. However, it was later showed this Zn^2+^-finger primarily stabilizes the dimerization loop and disrupting it does not impair dsDNA specificity ^11,26,34,39^. Here, we uncover an extended interaction network that tracks the B-form DNA architecture, spanning at least eight cGAS protomers across multiple 100-bp duplexes (Figures 3D and S5D-E). These interactions not only stabilize the minimal functional unit, but also drive assembly of the high-order protein•dsDNA mesh (Figures 3-4). We envision that single-stranded nucleic-acids would completely fail to support such a coordinated groove-tracking mechanism. Moreover, other double-helical nucleic acids would not be accommodated due to alternate helical architectures such as A-form dsRNA. Consistent with this notion, oligoadenylate synthases (OAS proteins), which share the NTase fold with cGAS, selectively recognize dsRNA ^48,49^. Structural comparison of cGAS•dsDNA and OAS1•dsRNA reveals that Arg^161^ of cGAS does not align with the minor groove of the A-form helix (Figure S7C). Moreover, OAS proteins lack both the junction-loops and N-terminal extension present in cGAS (Figure S7C), suggesting that these regions have diverged evolutionarily to differentiate between dsDNA- and dsRNA sensors. Intriguingly, nucleosomes suppress the autoactivation of cGAS in the nucleus by binding to the junction loops ^50–53^. Thus, host chromatin not only sequesters cGAS but also directly prevents its activation by masking key structural elements required for dsDNA recognition.

### The dsDNA length-dependent activation and percolation of cGAS

A major outstanding question in DNA-sensing pathways is how and why higher-order assembly on “long” dsDNA is necessary to maximize cGAS activation. Prior studies suggested several indirect and speculative explanations, including avidity effect ^27^, modulation of dimerization probability ^26^, condensation-coupled activation (multivalency) ^29^, protection of bound dsDNA from nucleases ^38^, and restricted diffusion of nucleotide substrates and the dinucleotide intermediate ^34^. Our results directly resolve this long-standing conundrum by demonstrating that mesh construction enables cross-stabilization between protomers from distinct pseudo-tetramers, reinforcing a critical interaction required for the dsDNA-dependent allosteric activation of cGAS (Figure 3B and D). This highly interwoven network packs cGAS active-sites no more than 70 Å apart (Figure 5), which would also be advantageous for recapturing the intermediate to complete cyclization ^34^. Beyond its innate immune signaling function, GAS has been implicated in mechanical transactions of dsDNA, such as double-strand break repair and replication ^21,22,24^, and it was reported that dsDNA flexibility plays a major factor in regulating cGAS activation ^54^. We demonstrate here that mesh construction not only contracts dsDNA, but also provides the mechanical resilience and kinetic stability to the active signaling complex (Figure 4), thus providing a foundation for a new concept that cGAS can sense the mechanical states of DNA. Future single-molecule and mechanical studies are warranted to further elucidate the rheological and mechanical properties of cGAS condensates.

Combined with our previous study ^34^, our results suggest that percolation is exclusively reserved for the cognate ligand of cGAS (dsDNA), whereas binding to noncognate nucleic acids results in segragative phase-separation that fails to active its enzymatic activity. It would be interesting to determine how other innate immune sensors couple or decouple associative and segregative routes to form condensates and how these different material properties regulate their signaling functions.

## Supporting information

SupMaterials

## Acknowledgement

We thank Dr. Edward Twomey for his advice in cryo-EM data processing and Dr. Archit Garg for discussion. S.M.T is an NSF GRFP fellow. This work was supported by NIH grant R21 AR085266 to B.A. and J.S.; R35 GM145363, Johns Hopkins Synergy Award, and Jerome L. Greene Foundation Scholars Award to J.S.

## Author Contributions

S.W. and J.S. conceptualized the project. S.W. C.M.S., S.M.T. performed all experiments, analyzed data, and wrote the paper. B.A edited the paper and co-supervised S.M.T. J.S. supervised the overall project, analyzed data, and wrote the paper.

