## Supplementary material for "Protein•DNA mesh assembly drives dsDNA-specific and duplex length-dependent activation of cGAS": SupMaterials

#### Materials and Methods

All experiments were performed at least three times (biological replicates), and *p* values were calculated using the two-tailed *t-test* assuming unequal variances (Microsoft Excel). All error bars are obtained from calculating standard deviations (Microsoft Excel). Graphs are made using Kaleidagraph (Syngergy), structural figures were made using UCSF ChimeraX, Figures were organized using Adobe Illustrator.

**Construct design.** Full-length mouse (m)cGAS$^{(FL)}$ ( UniProt ID: Q8C6L5; Origene, Cat. MC217197) was engineered with an N-terminal Thr-Gly linker, a TEV cleavage site (ENLYFQG), an Asn-Ser-Ser-Ser-Gly linker, a maltose-binding protein (MBP), and a hexa-histidine tag ($His_6$) tag (Figures S2A). The construct was subcloned into a pET28b vector (Novagen) expressed in *E. coli* BL21(DE3) cells (NEB). Site-directed mutagenesis was performed by PCR on this mcGAS$^{FL}$ construct to generate the following variants K382E (dimerization mutant), R222E/K240E/R241E (JxN3), R222E/K240E/R241E/R244E (JxN4), R144E/K145E (N$^{mut}$), R144E/K145E/R443E (NC$^{mut}$), and R433E.

**Protein Expression and Purification.** Plasmids encoding wild-type and mutant cGAS were expressed in *E. coli* BL21(DE3) cells induced with 0.3 mM IPTG at 16°C for 18 hours. Cells were lysed by sonication in lysis buffer (20 mM phosphate buffer, pH 7.5, 500 mM NaCl, 40 mM imidazole) and proteins were purified by immobilized metal ion affinity chromatography (HisTrap HP, GE Healthcare). For untagged cGAS, the fusion protein was cleaved by the TEV protease at 4°C overnight and the cleaved tag was removed sequentially using a nickel column followed by amylose resin. Untagged or fusion proteins were further purified by size-exclusion chromatography on a Superdex 200 pg column (Cytiva, 28989335) equilibrated in storage buffer (20 mM Tris, pH 7.5, 150 mM NaCl, 0.5mM TCEP, 10% glycerol). Purified cGAS proteins were concentrated to >30 mg/mL using centrifugal concentrators (Millipore Sigma, UFC9030), flash-frozen in liquid nitrogen, and stored at −80°C (see also [1]).

**Pyrophosphatase-coupled cGAS activity assay.** The nucelotidyl transferase (NTase) activity of cGAS was measured using pyrophosphatase-coupled assay as previously described [1-4]. Briefly, 100 nM cGAS was incubated with 50 nM of *E. coli* pyrophosphatase, 200 µM of a 1:1 mixure of ATP and GTP, and 400 nM dsDNA (binding sites normalized to 18-bp) in 85 µl reaction buffer (25 mM Tris-acetate pH 7.4, 125 mM potassium acetate pH 7.4, 1 mM TCEP, 5 mM $Mg(Ac)_2$, and 5% glycerol) at 25 ± 2°C. At each time point, four aliquots (20 µl each) were withdrawn and quenched with an equal volume of quench buffer (reaction buffer supplemented 25 mM EDTA) in a 384-well plate (Corning, 3640). Quenched samples were mixed with 10 µl malachite green color development solution and incubated for 45 min at 25 ± 2°C. Absorbance was measured at ~620 nm and compared with a standard curve of inorganic phosphate to determine phosphate concentrations. Background values from control reactions lacking recombinant cGAS were subtracted from all measurements. Apparent catalytic rates were determined from the slopes of control-subtracted phosphate production over time.

**Fluorescence-anisotropy binding assays.** Increasing concentrations of cGAS were added to reactions containing fluorescein-amidite (FAM)-labeled dsDNA (18-bp, 60-bp, and 100-bp; 5 nM final). Fluorescence anisotropy (FA) was measured on a Tecan M1000 plate reader as previously described [1-4]. Changes in FA were plotted against cGAS concentration and fit to the Hill equation.

For off-rate measurements, FAM-labeled dsDNA fragments (24-bp, 60-bp, and 100-bp; 5 nM final) were pre-incubated with each cGAS construct (600 nM). After 30 min incubation, 20 µM unlabeled $dsDNA_{60}$ (binding site normalized; 18-bp footprint) was added, and the decrease in FA was monitored over 30 min at 25 ± 2 °C using a Tecan M1000 plate reader (instrument deadtime: ~ 30 sec). The fraction bound was normalized to the FA value of each FAM-labeled dsDNA-cGAS complex before addition of unlabeled $dsDNA_{60}$. Time-dependent decreases in FA were plotted and fitted to a single-component exponential decay model. All dsDNA sequences were derived from the Herpesvirus (HSV) genome. Sense strand: 5'-TAA GAC ACG ATG CGA TAA ($dsDNA_{18}$ ends here) AAT CTG ($dsDNA_{24}$ ends here) TTT GTA AAA TTT ATT AAG GGT ACA AAT TGC CCT AGC ($dsDNA_{60}$ ends here) ACA GGG GTG GGG TTA GGG CCG GGT CCC CAC ACC CAA ACG C ($dsDNA_{100}$ ends here)-3'. FAM was attached to the 5' end of the sense strand.

**Cell Culture and Imaging.** cGAS variants were cloned into a pCMV6 vector containing a C-terminal mCherry tag. An empty pCMV6-mCherry vector was used as a negative control. HEK293T cells (ATCC, CRL-11268), authenticated by short tandem repeat (STR) profiling and confirmed to be mycoplasma-free, were used for transfection. Cells were seeded into a 12-well plate ($0.1\times10^6$ cells per well) containing 20-mm round coverslips. Mouse cGAS or vector plasmids (300 ng) were transfected at ~70% cell confluence using Lipofectamine 2000 (Invivogen). After 16 h, cells were washed twice with 1× PBS, fixed with 4% paraformaldehyde, and mounted on glass slides. Fluorescence mages were taken using a BioTek Cytation 5 Cell Imaging Multimode Reader and analyzed for puncta formation using BioTek Gen 5 software (Agilent).

**Measuring cGAS activity in HEK293T cells using dual-luciferase reporter.** HEK293T cells (ATCC, CRL-11268) were maintained in high-glucose DMEM medium (ThermoFisher) supplemented with 10% FBS at 37 °C and 5% $CO_2$. Cells were seeded in a 24-well plate ($6 \times 10^4$ cells per well) and incubated overnight. Plasmids (50 ng each) encoding empty vector or full-length cGAS variants were transfected using Lipofectamine 3000 (Invitrogen), together with 5 ng of Renilla luciferase plasmid, 10 ng of a plasmid encoding human STING, and 10 ng of a Firefly luciferase reporter driven by the IFN-Iβ promoter. After 24 h, cells were washed once with PBS and lysed in passive lysis buffer (Promega). Lysates were transferred to a white 96-well plate with a solid white flat bottom (Corning 3917) and analyzed for Firefly and Renilla Luciferase activity using the Dual-Luciferase Reporter Assay System (Promega) on a Synergy H1 plate reader equipped with a dual injector (BioTek). Firefly luciferase activity was normalized to Renilla luciferase and subsequently normalized to the empty vector control as described in [1].

**FRAP assays using labeled nucleic acids.** cGAS proteins were incubated with the indicated labeled dsDNA for 30 minutes. A series of prebleach images were taken to establish a baseline fluorescence followed by bleaching with 100% laser power for ~20 s. Images were then collected at indicated time, then processed and analyzed in ImageJ as described previously [5].

**cGAS phase separation assays.** Cellvis-384 Glass Bottom plates were cleaned with a 5% Hellmanex Solution, rinsed with deionized water (DI), and subsequently etched with 1 M KOH. The plates were then thoroughly rinsed with DI water, dried overnight, and sealed with aluminum foil. Prior to use, wells were unsealed and rinsed with reaction buffer (25 mM Tris acetate pH 7.4, 125 mM potassium acetate pH 7.4, 1 mM TCEP, 5 mM Mg $(Ac)_2$ at pH 7.4, 1 mg/ml bovine serum albumin (BSA), and 5% glycerol). Recombinant mouse cGAS (3 μM) was mixed with nucleic acids (3 μM; binding sites normalized to 18-bp; 5% Alexa Fluo 488-labelled) in a final volume of 25 μl. Images were acquired every 5 min using a Cytation 5 imaging plate reader (Agilent). Images were analyzed for condensate formation using the BioTek Gen5 software (Agilent).

**Cryo-EM sample preparation and data acquisition.** For the cGAS•dsDNA$_{66}$ complex, purified full-length fusion mouse cGAS (His$_6$–MBP–TEV–mcGAS; 200 μl at 370 μM, ~37.4 mg/ml) was mixed with 66-bp dsDNA (37 μl at 500 μM) and adjusted to a final volume of 800 μl using complex buffer (20 mM Tris, pH 7.5, 150 mM NaCl, 0.5 mM TCEP). For the cGAS•dsDNA100 complex, purified MBP-tagged mcGAS$^{FL}$ (100 μl at 370 μM) was combined with 100-bp dsDNA (9.4 μl at 500 μM) and brought to a final volume of 900 μl using the same complex buffer. Mixtures were incubated on ice for 30 minutes and subsequently loaded onto a Superdex 200 pg column (Cytiva, 28989335; equilibrated with complex buffer) to remove unbound cGAS. Each complex eluted as a single peak near the void volume (Figures S2-S4).

dsDNA$_{66}$ and dsDNA$_{100}$was prepared by annealing an equimolar amount of Forward and Reverse oligos (derived from mouse cGAS) at 95°C for 10 minutes. The sequence is 5'- CCG GAC AAG CTA AAG AAG GTG CTG GAC AAA TTG AGA TTG AAA CGC AAA GAT ATC TCG GAG GCG GCC (dsDNA66 stop here) GAG ACG GTG AAT AAA GTT GTG GAA CGC CTG CTG C - 3'

The peak fractions were used for Cryo-EM grid preparation. Briefly, 4 µL of either the cGAS-dsDNA66 complex (~ 2 mg/mL cGAS) or the cGAS-dsDNA100 complex (~1.2 mg/mL cGAS) was applied to Quantifoil® Holey Carbon R 2/2 300-mesh gold grids (Electron Microscopy Sciences, Q3100AR2) that had been glow-discharged using a PELCO easiGlow system (TED Pella, 91000) under the following conditions: plasma current 15 mA, process Timer 60 s, vacuum 0.37 mBar, preprocess hold 10 s, and negative polarity. After 30 s wait time, grids were blotted (blot force 2 and blot time 2s) and plunged-frozen in liquid ethane using a Vitrobot Mark IV (ThermoFisher Scientific) at 4 °C and 100% humidity.

Frozen grids were imaged on a ThermoFisher Titan Krios microscope equipped with a Falcon 4i camera with a direct electron detector and a Selectris energy filter (10-eV slit width) at the Johns Hopkins Beckman cryo-EM Center. Movies were collected in counting mode as 40-frame exposures at a pixel size of 0.93 Å/pixel with a total electron dose of 40 $e^{-}/Å^{2}$. Automated data collections were performed using EPU (ThermoFisher Scientific) with a beam-image shift acquisition scheme and a defocus range of −1.2 to −1.8 µm. A total of 25,781 movies were collected in one session for cGAS•$dsDNA_{66}$ complex, and 25,299 movies were collected for the cGAS•$dsDNA_{100}$ complex.

**Cryo-EM Data Processing**.

**cGAS•$dsDNA_{66}$ complex** (Figure S2). All Cryo-EM data were processed on CryoSPARC (v.4.5.3). For cGAS•$dsDNA_{66}$ complex, 25,781 EER movies were imported and subjected to patch motion correction and patch CTF estimation. An initial set of 7,519,001 particles were auto-picked using Blob Picker and extracted with a 400-pixel box. High-quality 2D classes were then used as templates for template-based particle picking, yielding 8,865,317 particles. Two rounds of 2D classifications were performed to remove junk particles, resulting in 1,013,271 particles for *ab-initio* reconstruction and heterogeneous refinement. Four rounds of heterogeneous refinements using five initial volumes produced a subset of 658,714 particles (96.97%), which was used for homogeneous refinement (3.25 Å), followed by non-uniform refinement (3.16 Å) and local refinement to generate the consensus map (3.12 Å). To improve local features, each cGAS dimer within the pseudo-tetramer were individually masked and refined, yielding local maps at 2.72 Å (dimer #1) and 2.79 Å (dimer #2). Conformational variability within the pseudo-tetramer was assessed using 3D Variability Analysis (3DVA) applied to the 3.12 Å consensus particle stack (658,714 particles). Variability was visualized in simple-mode movies, and further analyzed in the cluster mode to generate six clusters (maps), each map was subsequently refined using the non-uniform refinement routine.

**cGAS•dsDNA$_{100}$ complex** (Figure S3). Processing for cGAS-dsDNA100 dataset followed similar procedures. A total of 25,299 EER movies were imported, motion corrected, and CTF estimated. Using the Blob Picker, 3,084,630 particles were auto-picked and extracted with a 576-pixel box. High-quality 2D classes resembling pseudo-tetramer were then used as templates for template-based picking, yielding 5,621,467 particles. After two rounds of 2D classifications, 1,242,362 particles were retained for ab-initio reconstruction and heterogeneous refinement. Four rounds of heterogeneous refinements using five initial volumes produced a subset of 567,079 particles (96.01%), which were carried for homogeneous refinement (3.23 Å), followed by non-uniform refinement (3.20 Å) and local refinement to generate the consensus map (3.17

Å). Local masking and refinement of the individual dimers yielded local maps for at 2.89 Å (cGAS dimer #1) and 2.91 Å (cGAS dimer #2). Conformational variability was examined using 3DVA on the 3.17 Å consensus dataset (567,079 particles). Variability was visualized in simple-mode movies, and further analyzed in the cluster mode to generate six clusters (maps), each map was subsequently refined using the non-uniform refinement routine.

**Two Pseudo-tetramers Bound to Three 100-bp dsDNA Fragments** (Figure S4). The high-order assembly containing two pseudo-tetramers bound to three 100-bp DNA fragments was also reconstructed from the same dataset comprising 25,299 EER movies. For this assembly,154,972 particles were auto-picked using Blob Picker and extracted with a 576-pixel box. High-quality 2D classes exhibiting multiple-tetramer features were used for template-based particle picking, yielding 1,984,376 particles. Two rounds of 2D classifications reduced the dataset to 572,247 particles for *ab-initio* reconstruction and heterogeneous refinement. Four rounds of heterogeneous refinements using five initial volumes produced 253,296 particles (97.20 %). These were refined using homogeneous refinement (5.23 Å), followed by non-uniform refinement (3.88 Å) and local refinement to generate the consensus map (3.76 Å). To improve local resolution, particularly for the less well-resolved pseudo-tetramer #1 within the dimer-of-tetramers assembly, each pseudo-tetramer was masked and refined individually, yielding local refined maps of 3.75 Å (pseudo-tetramer #1) and 3.59 Å (pseudo-tetramer #2).

**Model building, refinement and structural analysis.** Model building was initiated with a crystal structure for the mouse cGAS catalytic domain bound to 18-bp dsDNA (PDB ID: 7UUX)

[1] using UCSF Chimera (v.1.15). The fitted model was then used as an initial model for manual DNA building in Coot (v.0.9.6). The final models for all structures were refined with real-space refinement in Phenix (v.1.20.1). All visualizations were performed in ChimeraX. Information for the deposited map and modeling is summarized in Extended Data Table 1.

**Data availability**. The cryo-EM reconstructions are deposited into the Electron Microscopy Data Bank (EMDB). For pseudo-tetramer structures, the consensus maps are the primary cryo-EM maps in each deposition and each composite map, local map, as applicable, and half maps are supplied as supplemental files in each deposition. For dimer of pseudo-tetramer, the composite maps are the primary cryo-EM map, and consensus map, local maps and half maps are supplied as supplemental files in each deposition. All structural coordinates are deposited in the Protein Data Bank (PDB). The accession codes for each state (EMDB, PDB) are: cGAS-dsDNA$_{66}$ pseudo-tetramer (EMD-76008, 11SN), cGAS-dsDNA100 pseudo-tetramer (EMD-76078, 11VP), Two pseudo-tetramers bound to Three 100-bp dsDNA Fragments (EMD-76134, 11WN).

**Single-molecule experiments.** Experiments were performed using the Lumicks C-Trap. The Lumicks C2 microfluidics system was passivated using the following steps: 0.1% BSA in PBS flowed at 0.4 bar pressure for 30 mins, followed by flowing PBS at 0.4 bar pressure for 10 mins to rinse, followed by 0.5% Pluronics F-127 in PBS at 0.4 bar for 30 mins, and finished with a final rinse of PBS at 0.4 bar for 30 mins. 4.38 μm streptavidin coated polystyrene beads (Spherotech cat no. SVP-40-5) were captured in each trap and used to tether biotinylated λdsDNA (Lumicks). Experimental buffer contained 25 mM Tris acetate at pH 7.5, 125 mM potassium acetate, 5% glycerol, 5 mM $Mg(AC)_2$, and 2 mM TCEP.

In force clamp (FC) experiments, tethered λdsDNA moved to the protein chamber and incubated for 3 minutes with the force clamp engaged at 2 pN then force data was recorded. In force extension (FE) experiments, tethered λdsDNA was first stretched by moving one of the captured beads at a constant pre-adjusted pulling rate of 0.2 μm/s in the buffer only channel and the resulting force was recorded. The λdsDNA was then moved to a channel containing each cGAS construct (3 μM) and incubated for 10 min. Subsequently, the λdsDNA was moved to the buffer only channel and stretched using a constant pulling speed of 0.2 μm/s then the resulting force was recorded.

Fitting the FE curves of naked λdsDNA using an extensible worm-like-chain (WLC) model [6] indicated that its contour length is ~ 16.1 μm under our buffer conditions (Figure S7B), which is consistent with the pulled distance at 30 pN [6]. We then estimated the contour length of each cGAS•λdsDNA in complex using the pulled distance at 30 pN. The work performed during each pull was calculated by integrating the area under each FE curve (https://github.com/Cstalli4/cTrapAnalysis). The stiffness of each cGAS-bound complex was estimated by taking the slope of each FE curve between ~10-11 μm.

### Figure S1

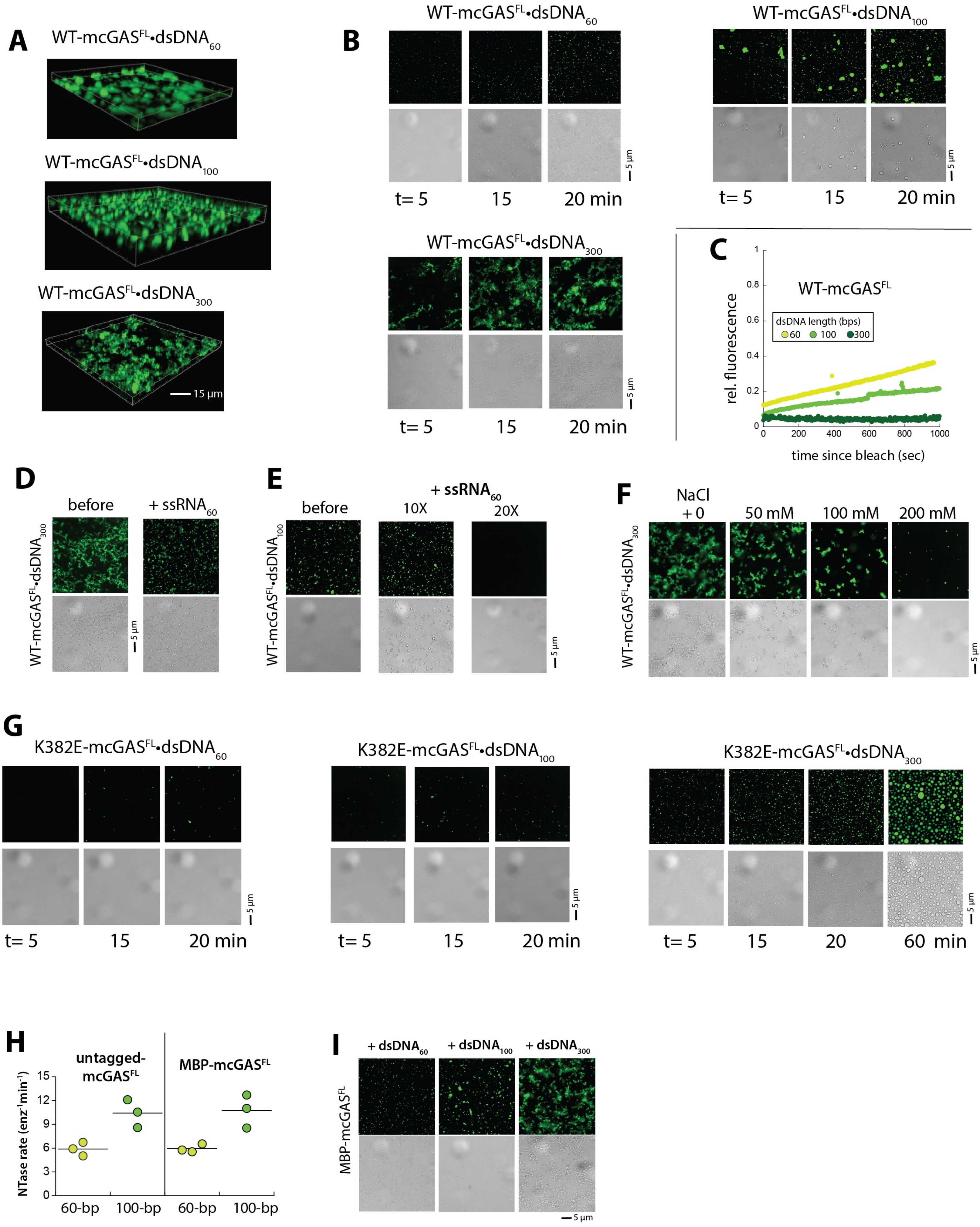
A
WT-mcGASFL•dsDNA60
WT-mcGASFL•dsDNA100
WT-mcGASFL•dsDNA300
15 µm
B
WT-mcGASFL•dsDNA60
WT-mcGASFL•dsDNA100
WT-mcGASFL•dsDNA300
5 µm
t= 5
15
20 min
C
WT-mcGASFL
dsDNA length (bps)
60
100
300
rel. fluorescence
time since bleach (sec)
D
before
+ ssRNA60
WT-mcGASFL•dsDNA300
E
+ ssRNA60
before
10X
20X
WT-mcGASFL•dsDNA100
F
NaCl
+ 0
50 mM
100 mM
200 mM
WT-mcGASFL•dsDNA300
G
K382E-mcGASFL•dsDNA60
K382E-mcGASFL•dsDNA100
K382E-mcGASFL•dsDNA300
t= 5
15
20
60 min
H
untagged-mcGASFL
MBP-mcGASFL
NTase rate (enz-1min-1)
60-bp
100-bp
I
+ dsDNA60
+ dsDNA100
+ dsDNA300
MBP-mcGASFL
5 µm

**Figure S1: Mouse cGAS forms hydrogel-like condensates via dimerization like the human enzyme.**

**(A)** Sample 3D images of WT-mcGAS$^{FL}$ condensates formed on different lengths of fluor-labeled dsDNA.

**(B)** Fluorescent and bright-field images of WT-mcGAS$^{FL}$ condensates formed on the fluor-labeled dsDNA of indicated length (3 μM for both protein and dsDNA).

**(C)** Another plot showing the recovery of fluorescence over time after bleaching for WT-mcGAS$^{FL}$ in complex with indicated length of dsDNA.

**(D)** Fluorescent and bright-field images of WT-mcGAS$^{FL}$•dsDNA$_{300}$ condensates (pre-incubated for 30 min) before and after incubating with unlabeled 60-base ssRNA in 20-fold access for 30 min.

**(E)** Fluorescent and bright-field images of WT-mcGAS$^{FL}$•dsDNA$_{100}$ condensates (pre-incubated for 30 min) before and after incubating with 60-base ssRNA in 10- and 20-fold access for 30 min. The lack of fluorescent signal indicates that dsDNA$_{100}$ has been completely displaced by 20-fold excess unlabeled ssRNA60 (condensates).

**(F)** Fluorescent and bright-field images of WT-mcGAS$^{FL}$•dsDNA$_{300}$ condensates (pre-incubated for 30 min) before and after incubating with increasing [NaCl] for 30 mins.

**(G)** Fluorescent and bright-field images of K382E-mcGAS$^{FL}$•dsDNAcondensates formed on dsDNA of different lengths.

**(H)** Catalytic activity of untagged and MBP-tagged mcGAS$^{FL}$ with different lengths of dsDNA (100 nM enzyme, 400 nM dsDNA, and 200 μM ATP/GTP)

**(I)** Fluorescent and bright-field images of MBP-mcGAS$^{FL}$•dsDNA condensates formed on dsDNA of different lengths (30 min incubation).

### Figure S2

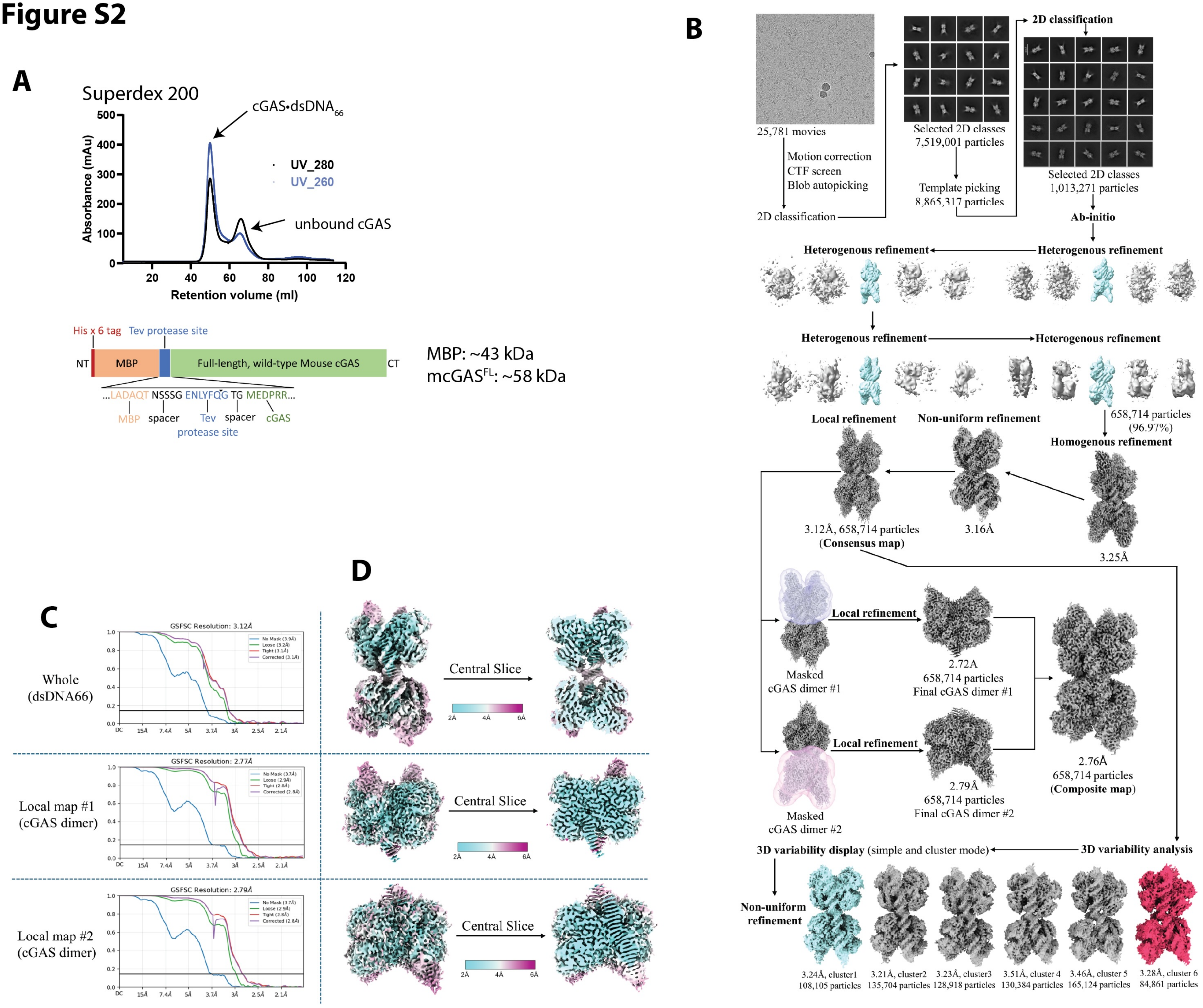

A
Superdex 200
cGAS•dsDNA66
UV_280
UV_260
unbound cGAS
Absorbance (mAu)
Retention volume (ml)
His x 6 tag
Tev protease site
NT
MBP
Full-length, wild-type Mouse cGAS
CT
...LADAQT NSSSG ENLYFQG TG MEDPRR...
MBP
spacer
Tev protease site
spacer
cGAS
MBP: ~43 kDa
mcGASFL: ~58 kDa
B
25,781 movies
Motion correction
CTF screen
Blob autopicking
2D classification
Selected 2D classes
7,519,001 particles
Template picking
8,865,317 particles
2D classification
Selected 2D classes
1,013,271 particles
Ab-initio
Heterogenous refinement
Heterogenous refinement
Heterogenous refinement
Heterogenous refinement
Local refinement
Non-uniform refinement
658,714 particles
(96.97%)
Homogenous refinement
3.12Å, 658,714 particles
(Consensus map)
3.16Å
3.25Å
Masked
cGAS dimer #1
Local refinement
2.72A
658,714 particles
Final cGAS dimer #1
Masked
cGAS dimer #2
Local refinement
2.79Å
658,714 particles
Final cGAS dimer #2
2.76Å
658,714 particles
(Composite map)
3D variability display (simple and cluster mode)
3D variability analysis
Non-uniform refinement
3.24Å, cluster1
108,105 particles
3.21Å, cluster2
135,704 particles
3.23Å, cluster3
128,918 particles
3.51Å, cluster 4
130,384 particles
3.46Å, cluster 5
165,124 particles
3.28Å, cluster 6
84,861 particles
C
Whole
(dsDNA66)
GSFSC Resolution: 3.12Å
Local map #1
(cGAS dimer)
GSFSC Resolution: 2.77Å
Local map #2
(cGAS dimer)
GSFSC Resolution: 2.79Å
D
Central Slice
Central Slice
Central Slice

**Figure S2: Cryo-EM workflow for the cGAS•$dsDNA_{66}$ pseudo-tetramer.**

**(A)** The size-exclusion chromatography (SEC) profile of MBP-tagged $mcGAS^{FL}$•$dsDNA_{66}$. The 260/280 ratio of the first peak indicates the presence of dsDNA. The second peak is without dsDNA as judged by the lower 260/280 ratio. A schematic for the recombinant $mcGAS^{FL}$ construct is also shown below.

**(B)** A flow chart showing cryo-EM data collection and refinement.

**(C)** FSC plots and **(D)** corresponding EM maps colored by resolution.

### Figure S3

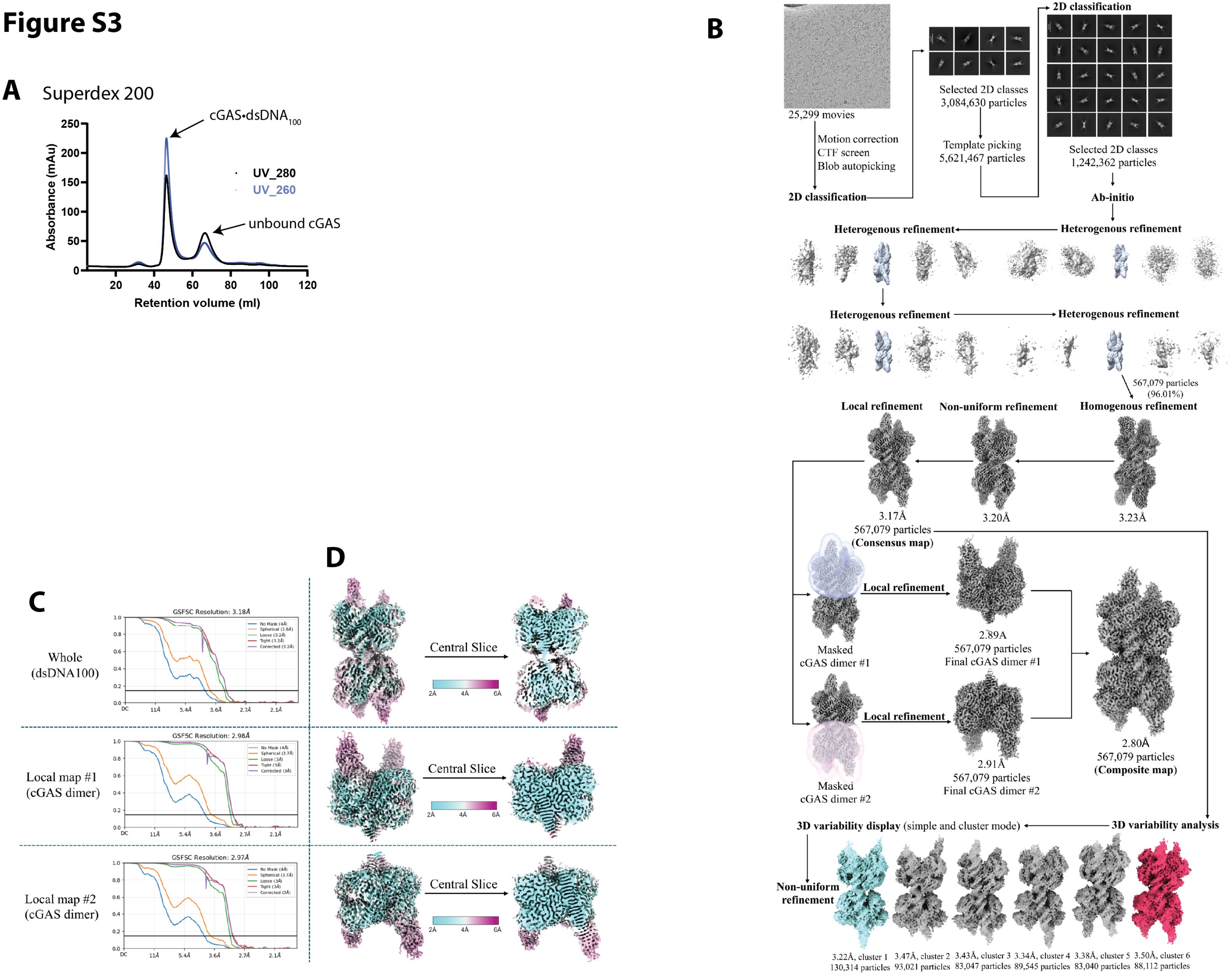
A
Superdex 200
cGAS•dsDNA100
unbound cGAS
UV_280
UV_260
Absorbance (mAu)
Retention volume (ml)
0
50
100
150
200
250
20
40
60
80
100
120
B
25,299 movies
Motion correction
CTF screen
Blob autopicking
2D classification
Selected 2D classes
3,084,630 particles
Template picking
5,621,467 particles
2D classification
Selected 2D classes
1,242,362 particles
Ab-initio
Heterogenous refinement
Heterogenous refinement
Heterogenous refinement
Heterogenous refinement
567,079 particles
(96.01%)
Local refinement
Non-uniform refinement
Homogenous refinement
3.17Å
567,079 particles
(Consensus map)
3.20Å
3.23Å
Masked
cGAS dimer #1
Local refinement
2.89A
567,079 particles
Final cGAS dimer #1
Masked
cGAS dimer #2
Local refinement
2.91Å
567,079 particles
Final cGAS dimer #2
2.80Å
567,079 particles
(Composite map)
3D variability display (simple and cluster mode)
3D variability analysis
Non-uniform
refinement
3.22Å, cluster 1
130,314 particles
3.47Å, cluster 2
93,021 particles
3.43Å, cluster 3
83,047 particles
3.34Å, cluster 4
89,545 particles
3.38Å, cluster 5
83,040 particles
3.50Å, cluster 6
88,112 particles
C
Whole
(dsDNA100)
GSFSC Resolution: 3.18Å
Local map #1
(cGAS dimer)
GSFSC Resolution: 2.98Å
Local map #2
(cGAS dimer)
GSFSC Resolution: 2.97Å
D
Central Slice
Central Slice
Central Slice
2Å
4Å
6Å

**Figure S3: Cryo-EM workflow for the cGAS•dsDNA$_{100}$ pseudo-tetramer.**

**(D)** The size-exclusion chromatography (SEC) profile of MBP-tagged mcGAS$^{FL}$•dsDNA$_{100}$. The 260/280 ratio of the first peak indicates the presence of dsDNA. The second peak is without dsDNA as judged by the lower 260/280 ratio.

**(E)** A flow chart showing cryo-EM data collection and refinement.

**(F)** FSC plots and **(D)** corresponding EM maps colored by resolution.

#### Figure S4

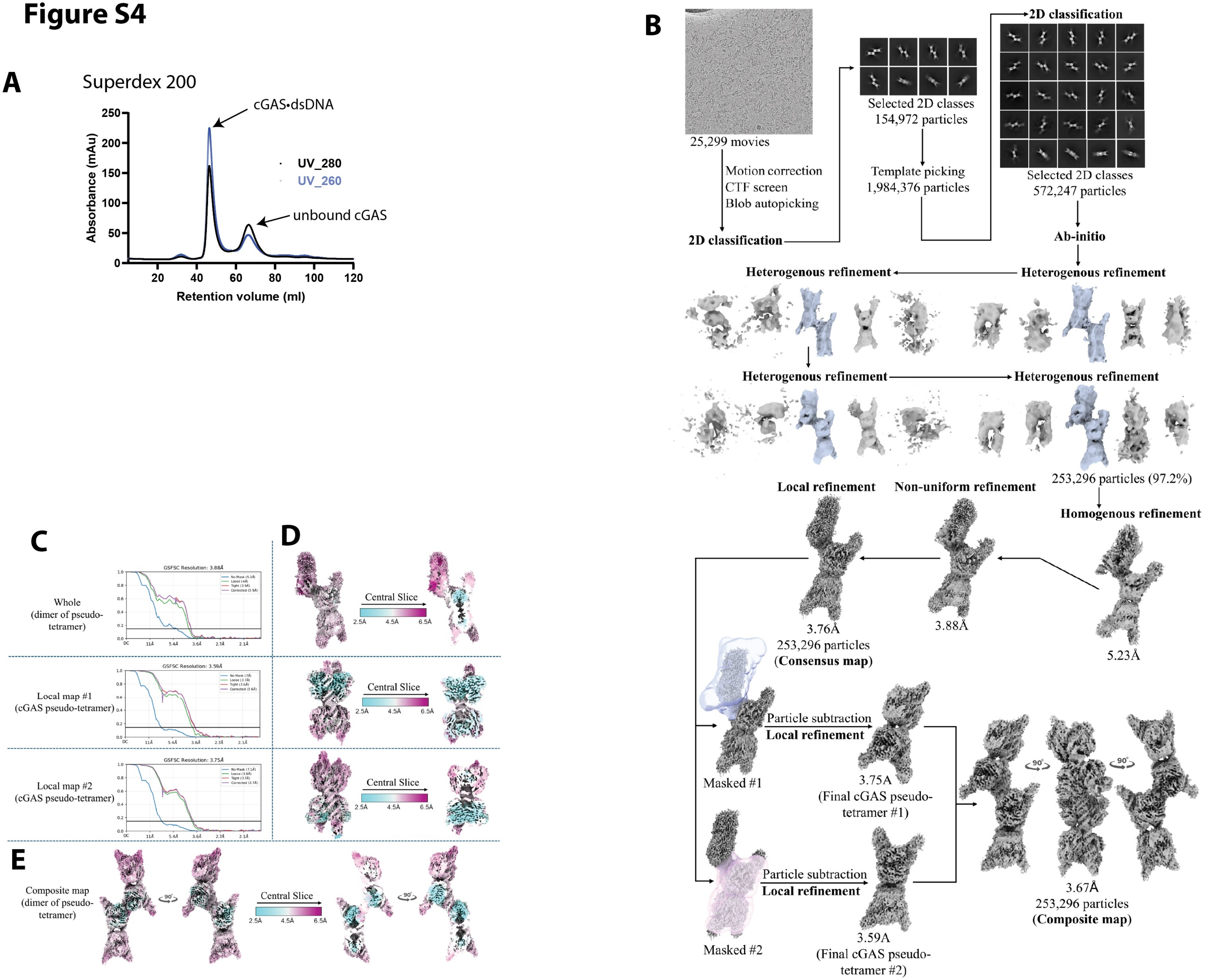

A
Superdex 200
cGAS•dsDNA
unbound cGAS
UV_280
UV_260
Absorbance (mAu)
Retention volume (ml)
B
25,299 movies
Motion correction
CTF screen
Blob autopicking
2D classification
Selected 2D classes
154,972 particles
Template picking
1,984,376 particles
2D classification
Selected 2D classes
572,247 particles
Ab-initio
Heterogenous refinement
Heterogenous refinement
Heterogenous refinement
Heterogenous refinement
253,296 particles (97.2%)
Homogenous refinement
Non-uniform refinement
Local refinement
5.23Å
3.88Å
3.76Å
253,296 particles
(Consensus map)
Masked #1
Particle subtraction
Local refinement
3.75A
(Final cGAS pseudo-tetramer #1)
Masked #2
Particle subtraction
Local refinement
3.59A
(Final cGAS pseudo-tetramer #2)
90°
90°
3.67Å
253,296 particles
(Composite map)
C
Whole
(dimer of pseudo-tetramer)
GSFSC Resolution: 3.88Å
Local map #1
(cGAS pseudo-tetramer)
GSFSC Resolution: 3.59Å
Local map #2
(cGAS pseudo-tetramer)
GSFSC Resolution: 3.75Å
D
Central Slice
2.5Å
4.5Å
6.5Å
E
Composite map
(dimer of pseudo-tetramer)
90°
Central Slice
2.5Å
4.5Å
6.5Å

**Figure S4: Cryo-EM workflow for the cGAS•dsDNA$_{100}$ dimer of pseudo-tetramers.**

**(A)** The size-exclusion chromatography (SEC) profile of MBP-tagged mcGAS$^{FL}$•dsDNA$_{100}$. The 260/280 ratio of the first peak indicates the presence of dsDNA. The second peak is without dsDNA as judged by the lower 260/280 ratio.

**(B)** A flow chart showing cryo-EM data collection and refinement.

**(C)** FSC plots and **(D-E)** corresponding EM maps colored by resolution.

### Figure S5

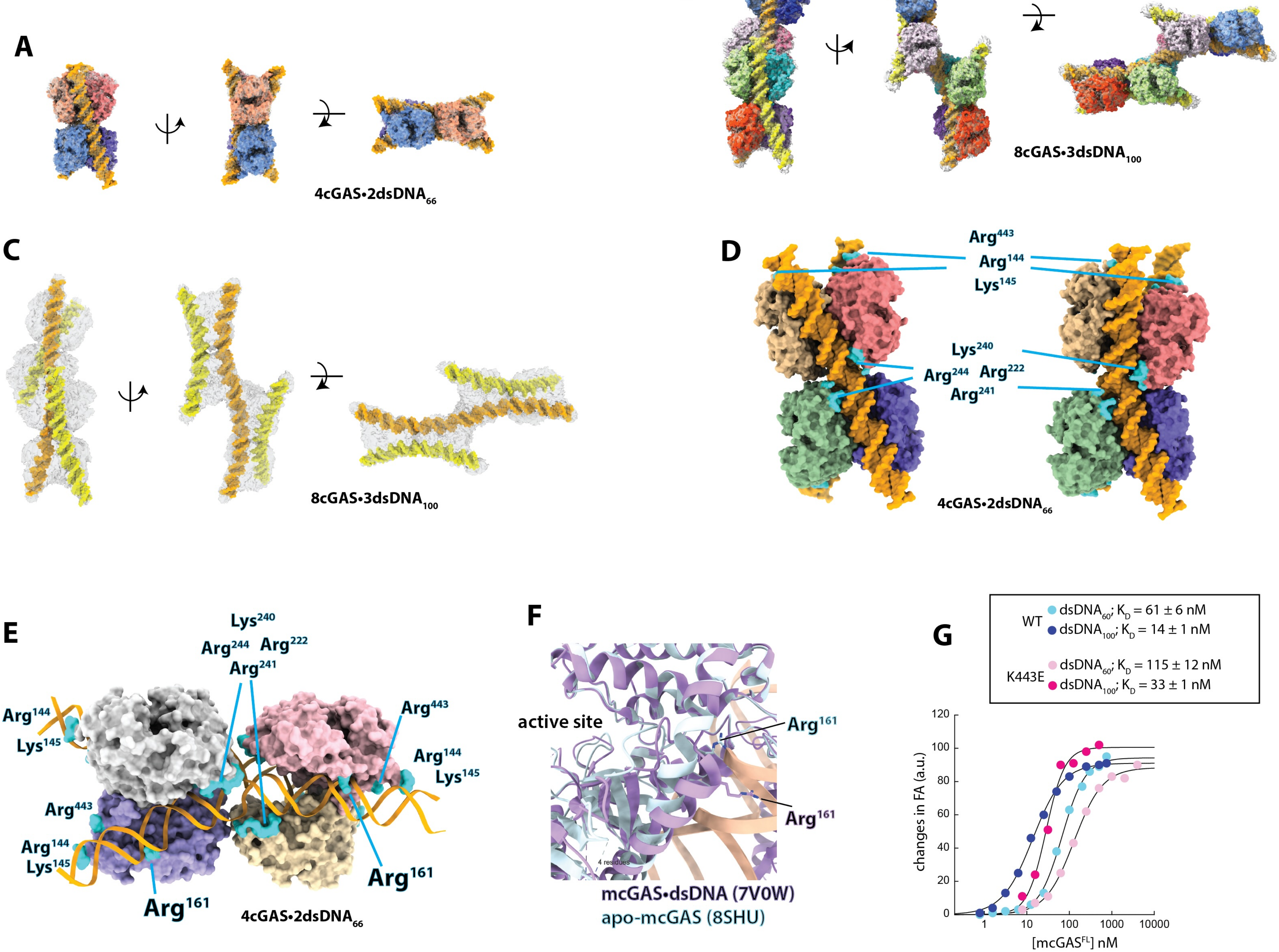
A
4cGAS•2dsDNA$_{66}$
B
8cGAS•3dsDNA$_{100}$
C
8cGAS•3dsDNA$_{100}$
D
Arg$^{443}$
Arg$^{144}$
Lys$^{145}$
Lys$^{240}$
Arg$^{244}$ Arg$^{222}$
Arg$^{241}$
4cGAS•2dsDNA$_{66}$
E
Lys$^{240}$
Arg$^{244}$ Arg$^{222}$
Arg$^{241}$
Arg$^{144}$
Lys$^{145}$
Arg$^{443}$
Arg$^{144}$
Lys$^{145}$
Arg$^{443}$
Arg$^{144}$
Lys$^{145}$
Arg$^{161}$
Arg$^{161}$
4cGAS•2dsDNA$_{66}$
F
active site
Arg$^{161}$
Arg$^{161}$
4 residues
mcGAS•dsDNA (7V0W)
apo-mcGAS (8SHU)
G
WT
dsDNA$_{60}$; $K_D$ = 61 ± 6 nM
dsDNA$_{100}$; $K_D$ = 14 ± 1 nM
K443E
dsDNA$_{60}$; $K_D$ = 115 ± 12 nM
dsDNA$_{100}$; $K_D$ = 33 ± 1 nM
changes in FA (a.u.)
[mcGAS$^{FL}$] nM
0
20
40
60
80
100
120
1
10
100
1000
10000

**Figure S5: Supporting cryo-EM and biochemical data.**

**(A)** The space filled model/map and additional views of the mcGAS$^{FL}$•dsDNA$_{66}$ pseudo-tetramer. Each dimer is colored in different shades of red and blue, respectively. dsDNA is colored in orange.

**(B)** The pace filled model/map and additional views of two mcGAS$^{FL}$ pseudo-tetramers formed on three dsDNA$_{100}$ fragments. Each monomer is colored differently. The shared dsDNA is colored in orange, and two peripheral DNA duplexes are colored in yellow.

**(C)** The space filled model/map and additional views 100-bp DNA duplexes from (**B**). The shared dsDNA in the middle is colored orange, and two peripheral DNA duplexes are colored in yellow.

**(D)** Two most distinct outputs of mcGAS$^{FL}$•dsDNA$_{66}$ from the 3DVA cluster mode are shown. Each monomer is colored differently, and the junction and N/C-terminal residues are colored in cyan.

**(E)** The cryo-EM structure of the mcGAS$^{FL}$•dsDNA$_{66}$ pseudo-tetramer. Residues that track either the major or minor groove of dsDNA are colored in cyan (same as those seen from the mcGAS$^{FL}$•dsDNA$_{100}$ structures in Fig. 3D).

**(F)** An overlay between the *apo*- and dsDNA-bound catalytic domain of mcGAS. Note the two different positions of Arg$^{161}$.

**(G)** Sample fluorescence anisotropy (FA) binding curves of WT- and R443E-mcGAS$^{FL}$ against FAM-labeled 60- or 100-bp dsDNA. $K_D$ values are listed. $n = 3$, ± standard deviations. a.u. = arbitrary units.

#### Figure S6

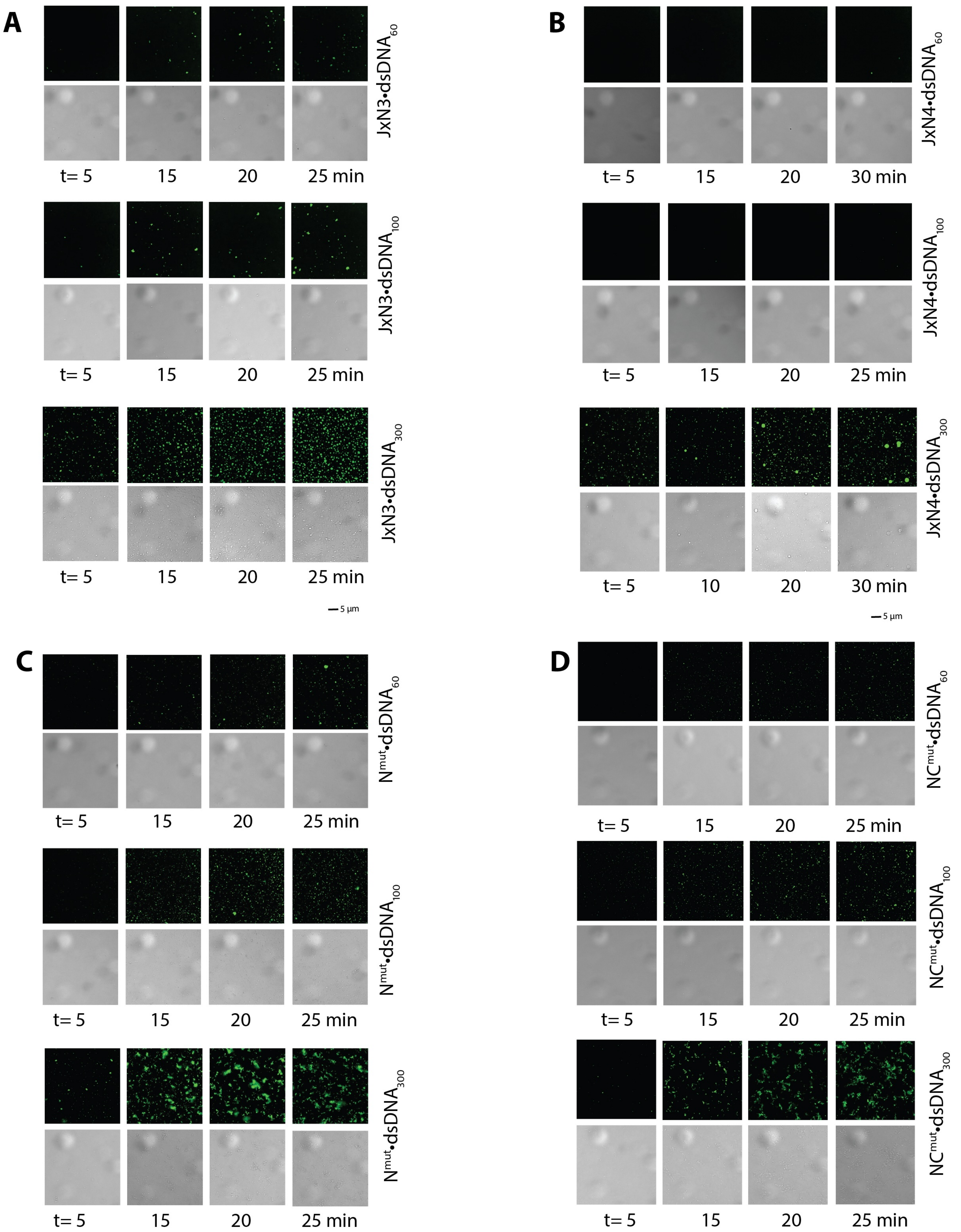

A
JxN3•dsDNA$_{60}$
t= 5
15
20
25 min
JxN3•dsDNA$_{100}$
t= 5
15
20
25 min
JxN3•dsDNA$_{300}$
t= 5
15
20
25 min
5 µm
B
JxN4•dsDNA$_{60}$
t= 5
15
20
30 min
JxN4•dsDNA$_{100}$
t= 5
15
20
25 min
JxN4•dsDNA$_{300}$
t= 5
10
20
30 min
5 µm
C
N$^{mut}$•dsDNA$_{60}$
t= 5
15
20
25 min
N$^{mut}$•dsDNA$_{100}$
t= 5
15
20
25 min
N$^{mut}$•dsDNA$_{300}$
t= 5
15
20
25 min
D
NC$^{mut}$•dsDNA$_{60}$
t= 5
15
20
25 min
NC$^{mut}$•dsDNA$_{100}$
t= 5
15
20
25 min
NC$^{mut}$•dsDNA$_{300}$
t= 5
15
20
25 min

**Figure S6: Fluorescent and bright field images of mcGAS$^{FL}$ mutant condensates formed on the dsDNA of indicated length.**

**(A)** JxN3.

**(B)** JxN4

**(C)** N$^{mut}$

**(D)** NC$^{mut}$

### Figure S7

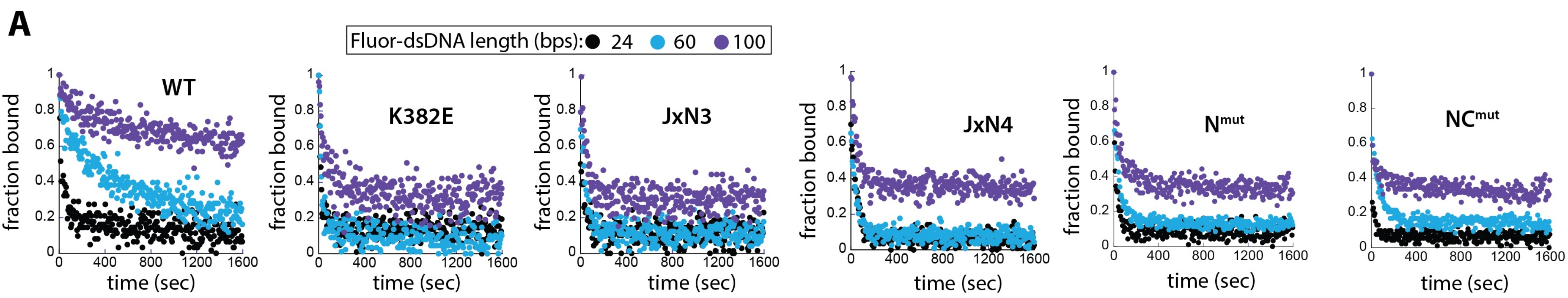

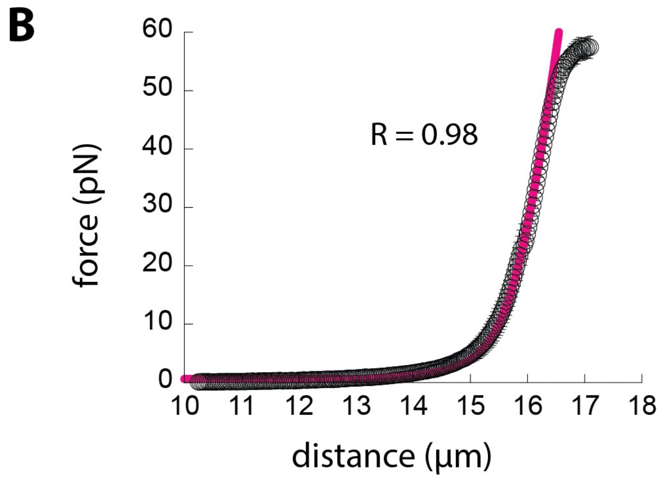

Contour length ($L_0$) = 16.1 ± 0.1 µm
Persistence length ($L_P$) = 50.0 ± 0.2 nm
Elastic Modulus ($K_0$) = 1330 ± 200 pN

$$\text{distance} = L_0 * (1-(1/2)*\text{sqrt}(k_BT/\text{force}*L_P)) + (\text{force}/K_0)$$

$k_B$ = boltzmann's constant
T = room temperature (25°C)

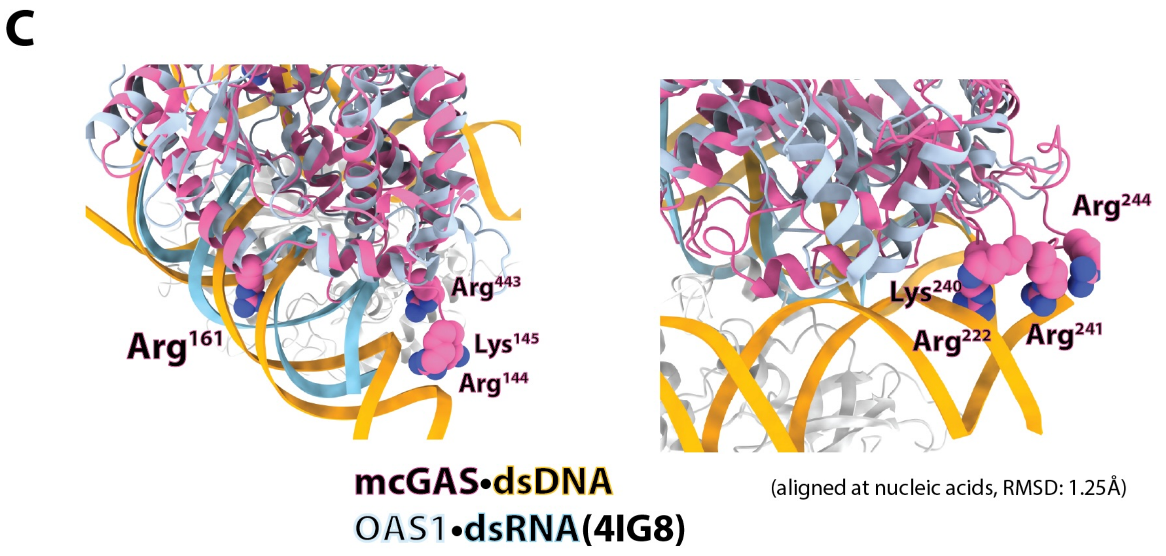

**Figure S7**

**(A)** Representative plots of off-rate kinetics experiment. Each mcGAS$^{FL}$ construct (600 nM) was incubated with the FAM-labeled dsDNA of indicated length (5 nM) for 30 min. The changes in fluorescence anisotropy (FA) upon adding excess dsDNA$_{60}$ (20 µM; unlabeled) was then tracked over time and the fraction remaining bound was calculated from the FA of the bound complex before adding dsDNA$_{60}$.

**(B)** The average FE curve of naked λdsDNA fit with an extensible worm-like-chain model (magenta line) [1]. The $L_0$, $L_P$, and $K_0$ values are averages of $n = 5$, ± standard deviation.

**(C)** An overlay between cGAS•dsDNA and OAS1•dsRNA.

#### Supplementary Table 1: Cryo-EM data collection, refinement, and validation statistics

| | cGAS•dsDNA66 (pseudo tetramer) (EMDB-76008) (PDB 11SN) | cGAS•dsDNA66 (local map #1) (EMDB-76004) (PDB 11SK) | cGAS•dsDNA66 (local map #2) (EMDB-76006) (PDB 11SL) |
|---|---|---|---|
| **Data collection and processing** | | | |
| Magnification | 130,000x | 130,000x | 130,000x |
| Voltage (kV) | 300 | 300 | 300 |
| Electron exposure ($e^{-}/Å^2$) | 40 | 40 | 40 |
| Defocus range (μm) | -1.2 to -1.8 | -1.2 to -1.8 | -1.2 to -1.8 |
| Pixel size (Å) | 0.93 | 0.93 | 0.93 |
| Symmetry imposed | C1 | C1 | C1 |
| Initial particle images (no.) | 8,865,317 | 8,865,317 | 8,865,317 |
| Final particle images (no.) | 658,714 | 658,714 | 658,714 |
| Map resolution (Å)<br>FSC threshold = 0.143 | 3.12 | 2.72 | 2.79 |
| Map resolution range (Å) | 2.5 – 38.1 | 2.0-42.9 | 2.3-43.7 |
| **Refinement** | | | |
| Initial model used (PDB code) | 7UUX | 7UUX | 7UUX |
| Model resolution (Å)<br>FSC threshold = 0.143 | 3.1 | 2.8 | 2.8 |
| Model resolution range (Å) | 3.0 – 3.3 | 2.7 – 3.0 | 2.7-3.1 |
| Map sharpening *B* factor ($Å^2$) | -89.7 | -97.4 | -90.7 |
| Model composition<br>Non-hydrogen atoms<br>Protein residues<br>Nucleotide<br>Ligands (Zn) | <br>16,156<br>1,456<br>200<br>4 | <br>8,365<br>728<br>114<br>2 | <br>8,324<br>728<br>112<br>2 |
| *B* factors ($Å^2$) (min/max/mean)<br>Protein<br>Nucleotide<br>Ligand (Zn) | <br>7.70/174.46/69.56<br>16.34/207.07/105.09<br>29.94/93.08/59.91 | <br>18.69/103.22/38.30<br>3.93/143.53/57.65<br>56.58/59.04/57.81 | <br>21.33/115.26/45.10<br>4.18/154.00/60.26<br>56.38/60.35/58.36 |
| R.m.s. deviations<br>Bond lengths (Å)<br>Bond angles (°) | <br>0.004<br>0.569 | <br>0.004<br>0.577 | <br>0.004<br>0.603 |
| Validation<br>MolProbity score<br>Clashscore<br>Poor rotamers (%) | <br>1.54<br>3.68<br>0.00 | <br>1.50<br>3.49<br>0.30 | <br>1.54<br>3.62<br>0.15 |
| Ramachandran plot<br>Favored (%)<br>Allowed (%)<br>Disallowed (%) | <br>94.48<br>5.52<br>0.00 | <br>94.75<br>5.25<br>0.00 | <br>94.34<br>5.66<br>0.00 |

| | cGAS-dsDNA100 (pseudo tetramer) (EMDB-76078) (PDB 11VP) | cGAS-dsDNA100 (local map #1) (EMDB-76076) (PDB 11VN) | cGAS-dsDNA100 (local map #2) (EMDB-76077) (PDB 11VO) |
|---|---|---|---|
| **Data collection and processing** | | | |
| Magnification | 130,000x | 130,000x | 130,000x |
| Voltage (kV) | 300 | 300 | 300 |
| Electron exposure ($e^{-}/Å^2$) | 40 | 40 | 40 |
| Defocus range (μm) | -1.2 to -1.8 | -1.2 to -1.8 | -1.2 to -1.8 |
| Pixel size (Å) | 0.93 | 0.93 | 0.93 |
| Symmetry imposed | C1 | C1 | C1 |
| Initial particle images (no.) | 5,621,467 | 5,621,467 | 5,621,467 |
| Final particle images (no.) | 567,079 | 567,079 | 567,079 |
| Map resolution (Å)<br>FSC threshold = 0.143 | 3.17 | 2.98 | 2.97 |
| Map resolution range (Å) | 2.0 – 33.8 | 2.4 – 48.1 | 2.4 – 47.3 |
| **Refinement** | | | |
| Initial model used (PDB code) | 7UUX | 7UUX | 7UUX |
| Model resolution (Å)<br>FSC threshold = 0.143 | 3.2 | 2.9 | 2.9 |
| Model resolution range (Å) | 3.1 – 3.5 | 2.8 – 3.1 | 2.8 – 3.1 |
| Map sharpening *B* factor ($Å^2$) | -93.1 | -99.8 | -98.1 |
| Model composition<br>Non-hydrogen atoms<br>Protein residues<br>Nucleotide<br>Ligands (Zn) | <br>16,156<br>1,456<br>200<br>4 | <br>7,996<br>728<br>96<br>2 | <br>8365<br>728<br>114<br>2 |
| *B* factors ($Å^2$) (min/max/mean)<br>Protein<br>Nucleotide<br>Ligand | <br>37.51/271.23/129.99<br>52.42/379.77/176.38<br>82.12/148.60/114.13 | <br>63.38/200.63/94.04<br>68.58/274.71/154.00<br>111.55/123.51/117.53 | <br>57.05/220.19/85.64<br>60.65/365.85/178.81<br>120.45/132.05/126.25 |
| R.m.s. deviations<br>Bond lengths (Å)<br>Bond angles (°) | <br>0.004<br>0.560 | <br>0.005<br>0.549 | <br>0.003<br>0.546 |
| Validation<br>MolProbity score<br>Clashscore<br>Poor rotamers (%) | <br>1.51<br>2.80<br>0.00 | <br>1.44<br>2.43<br>0.00 | <br>1.30<br>1.77<br>0.00 |
| Ramachandran plot<br>Favored (%)<br>Allowed (%)<br>Disallowed (%) | <br>93.16<br>6.84<br>0.00 | <br>93.78<br>6.22<br>0.00 | <br>94.75<br>5.25<br>0.00 |

| | cGAS-dsDNA100 (dimer of pseudo tetramer) (EMDB-76134) (PDB 11WN) | cGAS-dsDNA100 (local map #1) (EMDB-76131) (PDB 11WK) | cGAS-dsDNA100 (local map #2) (EMDB-76132) (PDB 11WL) |
|---|---|---|---|
| **Data collection and processing** | | | |
| Magnification | 130,000x | 130,000x | 130,000x |
| Voltage (kV) | 300 | 300 | 300 |
| Electron exposure ($e^{-}$/Å$^{2}$) | 40 | 40 | 40 |
| Defocus range (μm) | -1.2 to -1.8 | -1.2 to -1.8 | -1.2 to -1.8 |
| Pixel size (Å) | 0.93 | 0.93 | 0.93 |
| Symmetry imposed | C1 | C1 | C1 |
| Initial particle images (no.) | 1,984,376 | 1,984,376 | 1,984,376 |
| Final particle images (no.) | 253,296 | 253,296 | 253,296 |
| Map resolution (Å)<br>FSC threshold = 0.143 | 3.67 | 3.59 | 3.75 |
| Map resolution range (Å) | 3.2 – 58.6 | 2.9 – 39.5 | 3.0 – 50.5 |
| **Refinement** | | | |
| Initial model used (PDB code) | 7UUX | 7UUX | 7UUX |
| Model resolution (Å)<br>FSC threshold = 0.143 | 3.9 | 3.6 | 3.7 |
| Model resolution range (Å) | 3.6 – 4.9 | 3.5 – 4.0 | 3.6 – 4.7 |
| Map sharpening *B* factor (Å$^{2}$) | -81.75 | -82.4 | -81.1 |
| Model composition<br>Non-hydrogen atoms<br>Protein residues<br>Nucleotide<br>Ligands (Zn) | <br>31,656<br>2912<br>368<br>8 | <br>16,238<br>1,456<br>204<br>4 | <br>16,279<br>1,456<br>206<br>4 |
| *B* factors (Å$^{2}$) (min/max/mean)<br>Protein<br>Nucleotide<br>Ligand | <br>17.60/147.47/72.57<br>32.48/229.10/114.27<br>31.84/124.54/74.64 | <br>25.56/143.13/74.65<br>42.92/230.47/123.66<br>36.27/152.20/95.05 | <br>18.75/278.76/93.68<br>37.63/348.45/143.27<br>31.38/127.12/77.91 |
| R.m.s. deviations<br>Bond lengths (Å)<br>Bond angles (°) | <br>0.003<br>0.525 | <br>0.003<br>0.522 | <br>0.003<br>0.554 |
| Validation<br>MolProbity score<br>Clashscore<br>Poor rotamers (%) | <br>1.41<br>2.19<br>0.00 | <br>1.43<br>2.47<br>0.00 | <br>1.44<br>2.59<br>0.00 |
| Ramachandran plot<br>Favored (%)<br>Allowed (%)<br>Disallowed (%) | <br>93.82<br>6.18<br>0.00 | <br>94.06<br>5.94<br>0.00 | <br>94.07<br>5.93<br>0.00 |
